# NMDA receptor–BK channel coupling regulates synaptic plasticity in the barrel cortex

**DOI:** 10.1101/2020.12.30.424719

**Authors:** Ricardo Gómez, Laura E. Maglio, Alberto J. Gonzalez-Hernandez, Belinda Rivero-Pérez, David Bartolomé-Martín, Teresa Giraldez

## Abstract

Postsynaptic N-methyl-D-aspartate receptors (NMDARs) are crucial mediators of synaptic plasticity due to their ability to act as coincidence detectors of presynaptic and postsynaptic neuronal activity. However, NMDARs exist within the molecular context of a variety of postsynaptic signaling proteins, which can fine-tune their function. Here we describe a novel form of NMDAR suppression by large-conductance Ca^2+^- and voltage-gated K^+^ (BK) channels in the basal dendrites of a subset of barrel cortex layer 5 pyramidal neurons. We show that NMDAR activation increases intracellular Ca^2+^ in the vicinity of BK channels, thus activating K^+^ efflux and strong negative feedback inhibition. We further show that neurons exhibiting such NMDAR–BK coupling serve as high-pass filters for incoming synaptic inputs, precluding the induction of spike-timing-dependent plasticity. Together, these data suggest that NMDAR-localized BK channels regulate synaptic integration and provide input-specific synaptic diversity to a thalamocortical circuit.

## INTRODUCTION

Glutamate is the primary excitatory chemical transmitter in the mammalian central nervous system (CNS), where it is essential for neuronal viability, network function, and behavioral responses (Reiner and Levitz, 2018). Glutamate activates a variety of pre- and postsynaptic receptors, including ionotropic receptors (iGluRs) that form ligand-gated cation-permeable ion channels. The iGluR superfamily includes α-amino-3-hydroxy-5-methyl-4-isoxazolepropionic acid receptors (AMPARs), kainate receptors, and N-methyl-D-aspartate receptors (NMDARs), all of which form tetrameric assemblies that are expressed throughout the CNS (Traynelis et al., 2010).

NMDARs exhibit high sensitivity to glutamate (apparent EC_50_ in the micromolar range) and voltage-dependent block by Mg^2+^ (Mayer et al., 1984; Nowak et al., 1984), slow gating kinetics (Lester et al., 1990), and high permeability to Ca^2+^ (MacDermott et al., 1986; Mayer and Westbrook, 1987) (for a review, see (Paoletti et al., 2013)). Together, these characteristics confer postsynaptic NMDARs with the ability to detect and decode coincidental activity of pre- and postsynaptic neurons: presynaptic glutamate release bringing about occupation of the agonist-binding site and AMPAR-driven postsynaptic depolarization removing voltage-dependent Mg^2+^ block. The coincidence of these two events leads to NMDAR activation and Ca^2+^ influx through the channel (Caporale and Dan, 2008; Paoletti et al., 2013), which initiates several forms of synaptic plasticity (Malenka and Nicoll, 1993; Markram et al., 1997).

Large-conductance Ca^2+^- and voltage-gated K^+^ (BK) channels are opened by a combination of membrane depolarization and relatively high levels of intracellular Ca^2+^ (Kshatri et al., 2018a; Latorre et al., 2017). Such micromolar Ca^2+^ increases are usually restricted to the immediate vicinity of Ca^2+^ sources such as voltage-gated Ca^2+^ channels (VGCCs) (Marrion and Tavalin, 1998), ryanodine receptors (RyRs) (Chavis et al., 1998), and inositol 1,4,5-triphosphate receptors (InsP3Rs) (Zhao et al., 2010). However, Ca^2+^ influx through non-selective cation-permeable channels, including NMDARs, has also been shown to activate BK channels in granule cells from the olfactory bulb and dentate gyrus (Isaacson and Murphy, 2001; Zhang et al., 2018). In these neurons, Ca^2+^ entry through NMDARs opens BK channels in somatic and perisomatic regions, causing repolarization of the surrounding plasma membrane and subsequent closure of NMDARs. Because BK channel activation blunts NMDAR- mediated excitatory responses, it provides a negative feedback mechanism that modulates the excitability of these neurons (Isaacson and Murphy, 2001; Zhang et al., 2018). Thus, the same characteristics that make NMDARs key components in excitatory synaptic transmission and plasticity can paradoxically give rise to an inhibitory response when NMDARs are located in the proximity of BK channels. However, it is unclear whether functional NMDAR–BK coupling is relevant at dendrites and dendritic spines.

The barrel field area in the primary somatosensory cortex, also known as the barrel cortex (BC), processes information from peripheral sensory receptors for onward transmission to cortical and subcortical brain regions (Erzurumlu and Gaspar, 2020; Petersen, 2019). Sensory information is received in the barrel cortex from different nuclei of the thalamus. Among these nuclei, ventral posterior medial nucleus, ventrobasal nucleus, and posterior medial nucleus are known to directly innervate layer 5 pyramidal neurons (BC-L5PNs) (Agmon and Connors, 1992; Constantinople and Bruno, 2013; El-Boustani et al., 2020; Rodriguez-Moreno et al., 2020). In basal dendrites of BC-L5PN, coactivation of neighboring dendritic inputs can initiate NMDAR-mediated dendritically-restricted spikes characterized by large Ca^2+^ transients and long-lasting depolarizations (Nevian et al., 2007; Polsky et al., 2009; Schiller et al., 2000), providing the appropriate environment for BK activation.

To determine whether functional NMDAR–BK coupling plays a role in synaptic transmission, and potentially synaptic plasticity, we investigated the thalamocortical synapses at basal dendrites of BC-L5PNs. We found that suppression of NMDAR activity by BK channels occurs in the basal dendrites of about 40% of BC-L5PNs, where NMDAR activation triggers strong negative feedback inhibition by delivering Ca^2+^ to nearby BK channels. This inhibition regulates the amplitude of postsynaptic responses and increases the threshold for induction of synaptic plasticity. Our findings thus unveil a calibration mechanism that can decode the amount and frequency of afferent synaptic inputs by selectively attenuating synaptic plasticity and providing input-specific synaptic diversity to a thalamocortical circuit.

## RESULTS

### NMDAR activation opens BK channels in BC-L5PN basal dendrites

To investigate whether NMDA receptors are functionally coupled to other channel types in the basal dendrites of BC-L5PNs, we obtained whole-cell voltage-clamp recordings from pyramidal neurons located beneath layer 4 barrels in acute mouse brain slices (n = 108) (Figure 1A). Localization and large pyramidal-shaped somata suggest that we predominantly recorded from layer 5a neurons. In the presence of Mg^2+^-free ACSF supplemented with tetrodotoxin (TTX; 1 µM) and glycine (10 µM), puff application of 200 µM NMDA to the basal dendrites of BC-L5PNs evoked a NMDAR- resembling inward current in all neurons (Figure 1B). Remarkably, about 40% of the BC-L5PNs developed a slower outward current at holding potentials more positive than -40 mV (Figure 1B), which showed a clear dependence on membrane voltage (Figure 1C). These two populations of BC-L5PNs were assigned as A-type neurons (lacking the outward current; n = 65, 60.2%) and B-type neurons (showing the outward current; n = 43, 39.8%). Inward current amplitude was directly proportional to the holding potential in both populations (Figure 1D, left), indicating that a similar molecular species carries the inward current in both neuronal types. Interestingly, activation of the outward current in B-type neurons significantly reduced net inward current flow, decreasing the inward charge transfer (Figure 1D, right). When NMDA was applied at the same distance from the soma than previously but towards layer 4, only inward current was observed (Figure S1). These results suggest that NMDA can evoke an outward current in the basal dendrites of B-type BC-L5PNs, but not in the soma, oblique/truncated dendrites, or initial segment of the apical dendrite. Our findings are in contrast to what has been found in other neurons such as hippocampal CA1 pyramidal cells, where NMDAR-dependent outward currents were observed at the soma (Zorumski et al., 1989) but not at dendrites (Figure S2).

**Figure 1.**
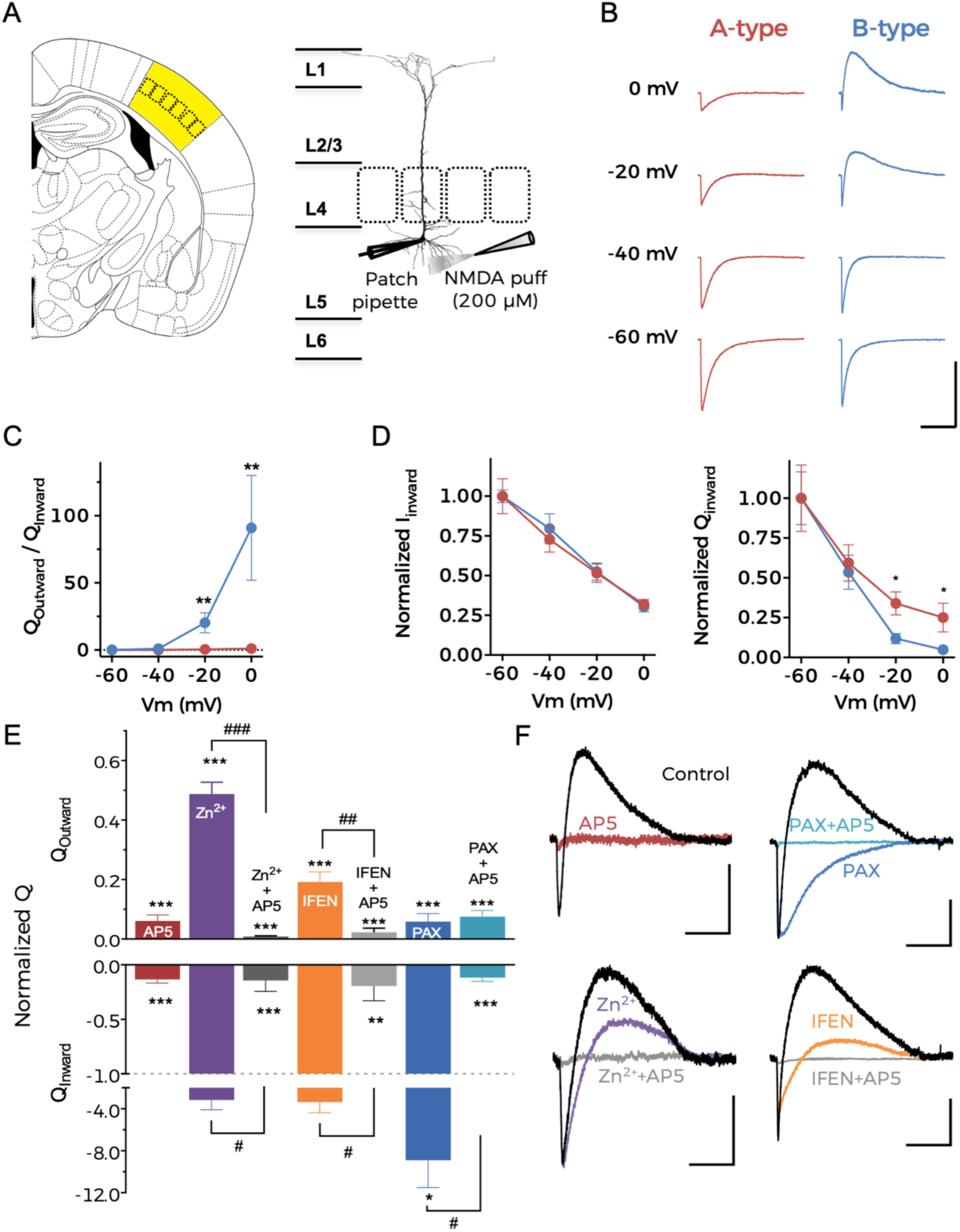
NMDAR activation opens BK channels in BC-L5PN basal dendrites. **(A)** Left, schematic representation of a mouse brain slice with the barrel cortex area highlighted in yellow. Right, schematic representation of the barrel cortex, depicting the BC-L5PNs under investigation. **(B)** Representative current traces obtained at the indicated holding potentials after application of NMDA to the basal dendrites. Scale bar represents 10 s and 200 pA. **(C)** Average charge-voltage (Q-V) relationships for A-type (red) and B-type (blue) neurons. **(D)** Normalized current-voltage (I-V, left) and Q-V (right) relationships for NMDAR inward currents from A-type and B-type neurons. Data points in **(C)** and **(D)** represent mean ± SEM. A-type: n = 15, B-type: n = 8; *p<0.05 and **p<0.01 (B-type vs. A-type). **(E)** Pharmacological characterization of NMDA-activated current in BC-L5PN basal dendrites. Normalized charge for the outward (Q_Outward_, top) and inward (Q_Inward_, bottom) component after the addition of different drugs. Data correspond to mean ± SEM. **p<0.01 and ***p<0.001 (treatment vs. ACSF); #p<0.05, ##p<0.01, and ###p<0.01 (drug+AP5 vs. drug alone). **(F)** Representative current traces obtained at -20 mV after NMDA application to the basal dendrites of B-type neurons in control conditions (ACSF, black traces) and after the application of different drugs. Traces are color-coded corresponding to the different treatments shown in **(E)**. Scale bars represent 5 s and 50 pA. In **(E)** and **(F)**, AP5, D-AP5 (100 µM; n = 6); IFEN, Ifenprodil (5 µM; n = 5); PAX, paxilline (1 µM; n = 5); Zn^2+^, ZnCl_2_ (100 nM; n = 5). See also Table S1.

We performed pharmacological characterization of inward and outward currents in B-type neurons at a holding potential of -20 mV (Figure 1E). The complete abrogation of inward current by the selective NMDAR antagonist D-AP5 (AP5; 100 µM) confirmed that it was due to cation flow through NMDARs. Moreover, because outward current was also abolished in the presence of AP5, we surmised that NMDAR activation is mandatory for outward current generation (Figure 1E and 1F). Selective inhibition of GluN2A- or GluN2B-containing NMDARs using 100 nM ZnCl_2_ or 5 µM ifenprodil, respectively, partially suppressed the outward current (Figures 1E and 1F). This suggests that inward current flow through both GluN2A- and GluN2B-containing NMDARs leads to activation of the outward current in B-type neurons.

The voltage dependence and direction of net ionic flow (Figure 1C) suggests that NMDAR-dependent outward current is driven by a voltage-dependent K^+^ channel. Because this current is reminiscent of that carried by BK channels in granule cells from the olfactory bulb and dentate gyrus (Isaacson and Murphy, 2001; Zhang et al., 2018), we applied NMDA to basal dendrites of BC-L5PNs in the presence of the specific BK channel pore blocker paxilline (1 µM). Paxilline completely abolished the NMDAR-dependent outward current without eliminating the inward component, suggesting that NMDAR activation in the basal dendrites of B-type neurons causes BK channels to open, and therefore that NMDARs and BK channels are functionally coupled.

### NMDARs and BK channels are within functional proximity in B-type BC-L5PNs

Our results suggested that activation of NMDARs in the basal dendrites of B-type BC-L5PNs provides both the membrane depolarization and Ca^2+^ needed to activate BK channels. To confirm this hypothesis, we recorded NMDA-evoked current in the presence of the intracellular Ca^2+^ chelators BAPTA and EGTA, which we delivered through the recording pipette. BAPTA and EGTA are useful biochemical tools to estimate the linear distance that Ca^2+^ diffuses from its source (Naraghi and Neher, 1997). Both chelators present similar affinities for Ca^2+^ but BAPTA has a faster association rate than EGTA, therefore outward current through BK channels would be recorded in the presence of EGTA, but not BAPTA, if both channels are located in close proximity. If the distance between both channels is larger, Ca^2+^ entering through NMDARs would be captured by both chelators before reaching BK, and no outward current would be observed.

We failed to observe NMDA-induced outward currents in any BC-L5PNs in the presence of 15 mM BAPTA (Figure 2A and 2D), confirming that Ca^2+^ entry through NMDARs is responsible for activation of BK channels. However, when the slower chelator EGTA was used at the same concentration, two populations of BC-L5PNs could be distinguished (Figure 2B). As in control conditions, the outward current in B-type neurons activated at holding potentials positive to -40 mV and exhibited a clear dependence on membrane voltage, but its amplitude was significantly reduced due to substantial Ca^2+^ chelation (compare Figure 2E with Figure 1C). When BAPTA concentration was reduced to 1 mM, outward current could again be recorded (Figure 2C) and was quantitatively identical to that obtained in control conditions (compare Figure 2F with Figure 1C). Using described Ca^2+^ linear diffusion data in the presence of BAPTA and EGTA (Naraghi and Neher, 1997), we estimated that NMDAR and BK channels are located within 15-60 nm of each other in the basal dendrites of B-type BC-L5PNs (Figure 2G); in close enough proximity to explain the specific functional coupling that we observed.

**Figure 2.**
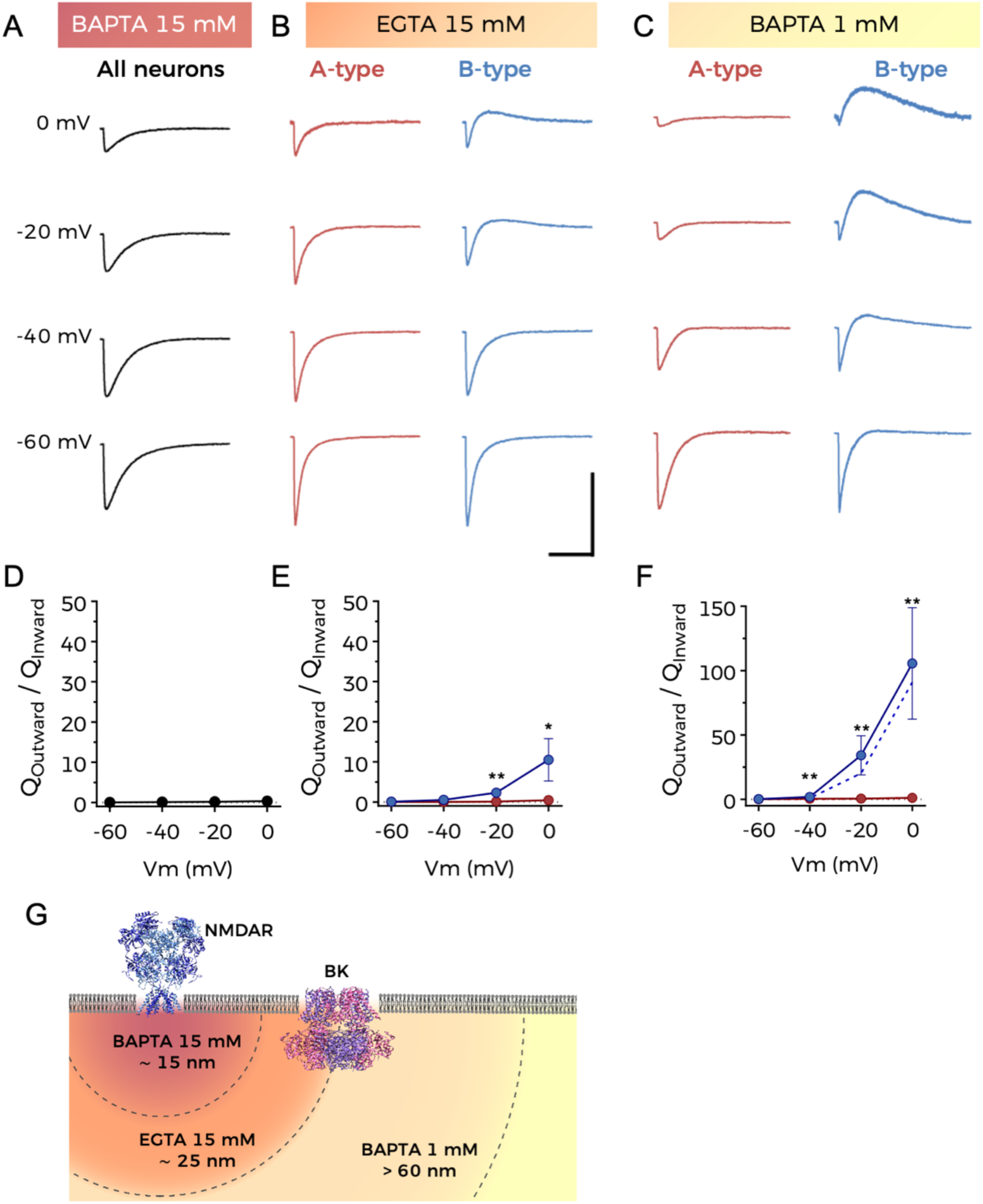
NMDARs and BK channels are within functional proximity in B-type BC-L5PNs. **(A–C)** Representative current traces obtained at the indicated holding potentials after application of NMDA to the basal dendrites of BC-L5PNs in the presence of 15 mM BAPTA **(A),** 15 mM EGTA **(B)**, and 1 mM BAPTA **(C)** in the recording pipette. Scale bars represent 10 s and 200 pA. **(D–F)** Average Q-V relationships corresponding to experiments described in **(A–C)**, respectively. Data points represent mean ± SEM; n = 18 **(D)**. Data points represent mean ± SEM; A-type: n = 8, B-type: n = 5 **(E)**. Data points represent mean ± SEM; A-type: n = 9, B-type: n = 5 **(F)**. Dashed lines represent data taken from Figure 1C for A-type **(E)** and B-type **(F)** neurons in the absence of Ca^2+^ chelators; *p<0.05 and **p<0.01 (B-type vs. A-type). **(G)** Schematic representation of the relative location of NMDARs and BK channels in the plasma membrane. Distances are estimated from the experimental data. See also Table S1.

### Both GluN2A- and GluN2B-containing NMDARs can functionally couple to BK channels

Because GluN2 subunits determine the deactivation kinetics and ion conductance of NMDARs (Vicini et al., 1998), we reasoned that the molecular composition of NMDARs might affect NMDAR–BK coupling efficiency. We first tested whether BK preferentially associates with specific NMDAR subunit combinations using a proximity ligation assay (PLA). In agreement with our pharmacological data (Figure 1F), positive PLA signals were observed for HEK293T cells co-expressing BK and either GluN1/GluN2A or GluN1/GluN2B NMDARs (Figure 3A), without any preference between GluN2A and GluN2B subunits (Figure 3B).

**Figure 3.**
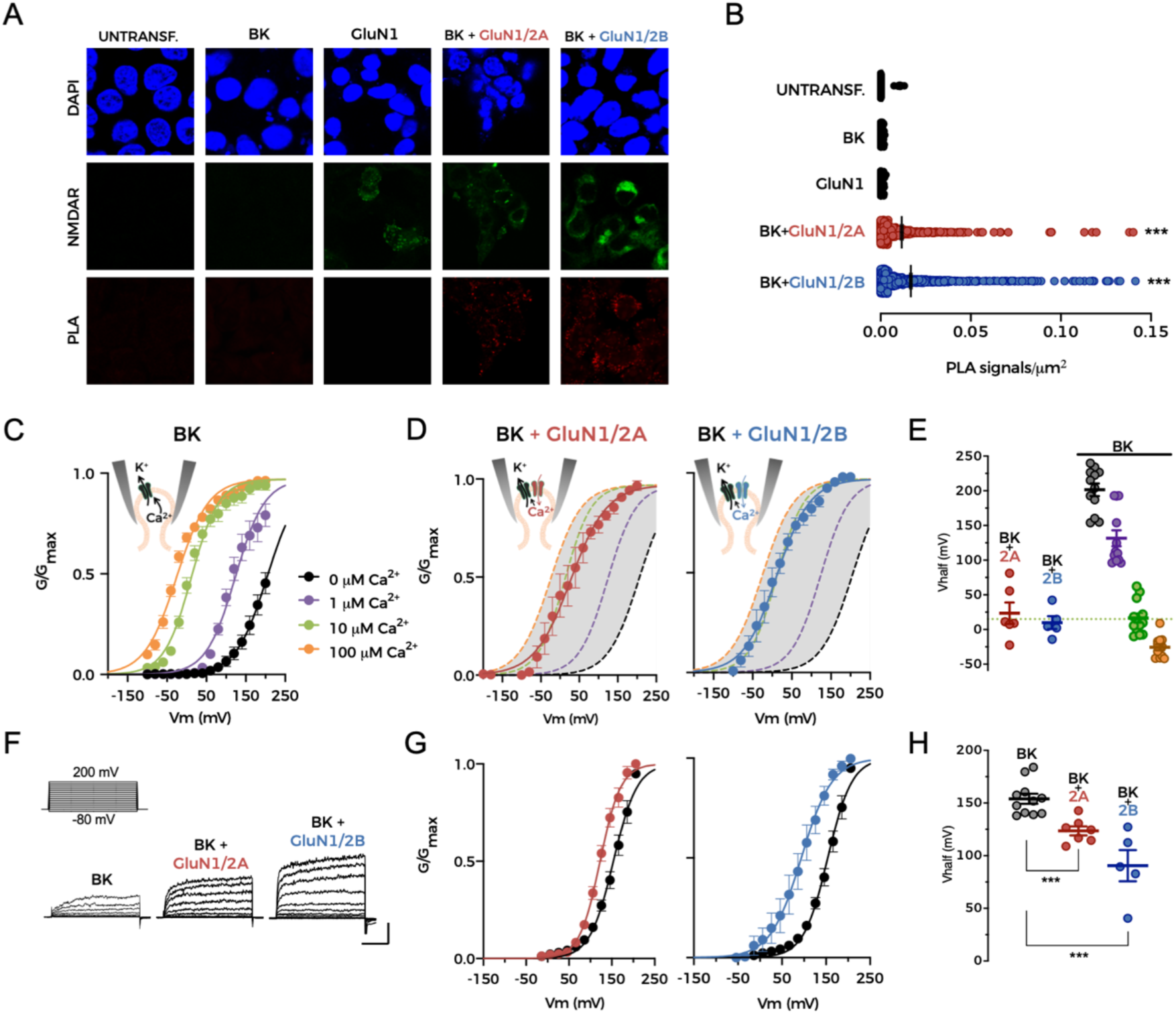
Both GluN2A- and GluN2B-containing NMDARs can functionally couple to BK channels. **(A)** Representative confocal microscopy images of PLA experiments in HEK293 cells expressing the protein combinations indicated at the top of the images. Each row corresponds to an imaging channel (top, DAPI, blue; middle, NMDAR, green; bottom, PLA, red). **(B)** Average PLA signals/µm^2^ for all conditions shown in **(A)**. Data points represent individual measurements, and each condition was studied in four independent experiments. Black lines represent mean ± SEM. ***p<0.001 vs. untransfected cells (UNTRANSF), BK alone (BK), and GluN1 alone (GluN1). **(C)** Normalized conductance-voltage (G-V) relationships obtained from patches expressing BK channels alone (n = 12) in the presence of different intracellular Ca^2+^ concentrations in symmetrical K^+^ solutions. Data points represent mean ± SEM. Lines represent the best fit of a Boltzmann equation to the data. Ca^2+^ concentrations are color-coded as indicated in the legend. **(D)** Normalized G-V relationships from patches co-expressing BK channels with GluN1/GluN2A (left; n = 6) or GluN1/GluN2B (right; n = 5) in the absence of intracellular Ca^2+^. Data points represent mean ± SEM. Lines represent the best fit of a Boltzmann equation to the data. **(E)** V_half_ values obtained from experiments depicted in **(C)** and **(D)**. Data points represent individual measurements and lines represent mean ± SEM. **(F)** Representative current traces recorded in physiological solutions from HEK293 cells expressing BK alone (left), BK+GluN1/GluN2A (middle), and BK+GluN1/GluN2B (right). Scale bars represent 30 ms and 2000 pA. **(G)** Normalized G-V relationships for BK channels co-expressed with GluN1/GluN2A (left, red; n = 6) or GluN1/GluN2B (right, blue; n = 5) in the absence of intracellular Ca^2+^. Black traces in both graphs correspond to G-V curves for BK channels expressed alone (n = 11). Data points represent mean ± SEM. Lines represent the best fit of a Boltzmann equation to the data. **(H)** Summary of V_half_ values from experiments in **(G)**. Data points represent individual measurements and lines represent mean ± SEM. ***p<0.001 vs. BK alone. See also Table S1.

Having confirmed that NMDARs and BK channels are located in close proximity in the plasma membrane when co-expressed in HEK293T cells, we characterized functional coupling between specific channel/subunit combinations. By using the inside-out configuration of the patch-clamp technique, we were able to monitor channel function while controlling ‘intracellular’ Ca^2+^ concentration in the bath solution (Giraldez et al., 2005). NMDARs were activated by including 200 µM NMDA and 10 µM glycine in the ‘extracellular’ pipette solution. The relative conductance (G) of BK channels exposed to different intracellular Ca^2+^ concentrations (from 0 to 100 µM) in patches from cells expressing BK channels but no NMDARs (Figure 3C) corresponded to typical Ca^2+^- dependent activation curves for these channels (Horrigan and Aldrich, 2002; Kshatri et al., 2018b). We reasoned that addition of Ca^2+^ to the intracellular side of the patch through a Ca^2+^ source such as NMDARs would result in a leftward shift of the BK activation curve in zero Ca^2+^ bath solution. Indeed, co-expression of BK channels with either GluN1/GluN2A or GluN1/GluN2B NMDARs produced a significant leftward shift of the BK activation curve (Figure 3D). Interestingly, the shift resulted in an activation curve comparable to that recorded with 10 µM intracellular Ca^2+^ in patches expressing BK alone (Figure 3E and Table S1). These results suggest that NMDAR activation increases intracellular Ca^2+^ concentration in the vicinity of BK channels, favoring their activation.

NMDARs are non-selective cation channels that are permeable to Na^+^, K^+^, and Ca^2+^ (Paoletti et al., 2013; Traynelis et al., 2010). Under physiological conditions, the proportion of NMDAR current carried by Ca^2+^ corresponds to 10-15% of the total current (Burnashev et al., 1995; Garaschuk et al., 1996; Plant et al., 1997). However, because our inside-out patch experiments were performed in the absence of Na^+^, inward current through NMDARs could only be due to Ca^2+^ ions, whose permeability is increased as extracellular Na^+^ concentration is reduced (Mayer and Westbrook, 1987). We may therefore be overestimating the effects of NMDAR-dependent Ca^2+^ activation on BK channels in our zero Na^+^ experimental conditions. Nevertheless, activation of GluN1/GluN2A or GluN1/GluN2B NMDARs in physiological concentrations of Na^+^ still produced a leftward shift in the BK activation curve (Figures 3F–3H). Interestingly, GluN1/GluN2B produced larger shifts than GluN1/GluN2A.

Taken together, these results demonstrate that both GluN2A- and GluN2B-containing NMDARs can provide sufficient Ca^2+^ for BK activation, consistent with our data from BC-L5PN basal dendrites (Figure 1F).

### A subpopulation of regular-spiking BC-L5PNs exhibit NMDAR–BK functional coupling

BC neurons can be classified into fast-spiking nonpyramidal GABAergic interneurons and regular-spiking or intrinsically-bursting pyramidal neurons according to their electrophysiological properties (Agmon and Connors, 1989, 1992; McCormick et al., 1985). Regular-spiking neurons from layers 5 and 6, where regularly-timed trains of action potentials are observed in response to somatic current injection, can be further characterized by the presence or absence of a depolarizing afterpotential known as a Ca^2+^ spike (Agmon and Connors, 1992; Chagnac-Amitai and Connors, 1989). To determine whether NMDAR–BK functional coupling is restricted to a specific neuronal subtype, we performed whole-cell current-clamp recordings in regular-spiking BC-L5PNs in mouse brain slices (n = 197) in the presence of physiological concentrations of Mg^2+^ and Ca^2+^. By inducing single action potentials in BC-L5PNs, we confirmed the presence of two populations of neurons that either exhibited a Ca^2+^ spike (n = 130, 66.0%) or not (n = 67, 34.0%) (Figure 4A and 4B), similar to those previously described (Agmon and Connors, 1992; Chagnac-Amitai and Connors, 1989; Maglio et al., 2018). We next investigated whether these two populations of BC-L5PNs could be correlated with NMDAR–BK functional coupling by subsequently perfusing the brain slices with a Mg^2+^-free solution supplemented with TTX (1 µM) and glycine (10 µM). Voltage clamp recordings were carried out in the same neurons upon puff application of NMDA to their basal dendrites (Figure 4C). None of the BC-L5PNs exhibiting a Ca^2+^ spike presented an outward current in response to NMDAR activation. Conversely, all BC-L5PNs lacking a Ca^2+^ spike showed a robust BK outward current after NMDA application (Figures 4B and 4C). No further intrinsic or evoked differences were observed between A-type and B-type neurons (Figures 4D–4I), except for those that were directly related to the presence of the Ca^2+^ spike (Figures 4J and 4K). We therefore conclude that the presence of NMDAR–BK coupling in basal dendrites is restricted to a population of regular-spiking BC-L5PNs that are characterized by the absence of Ca^2+^ spikes.

**Figure 4.**
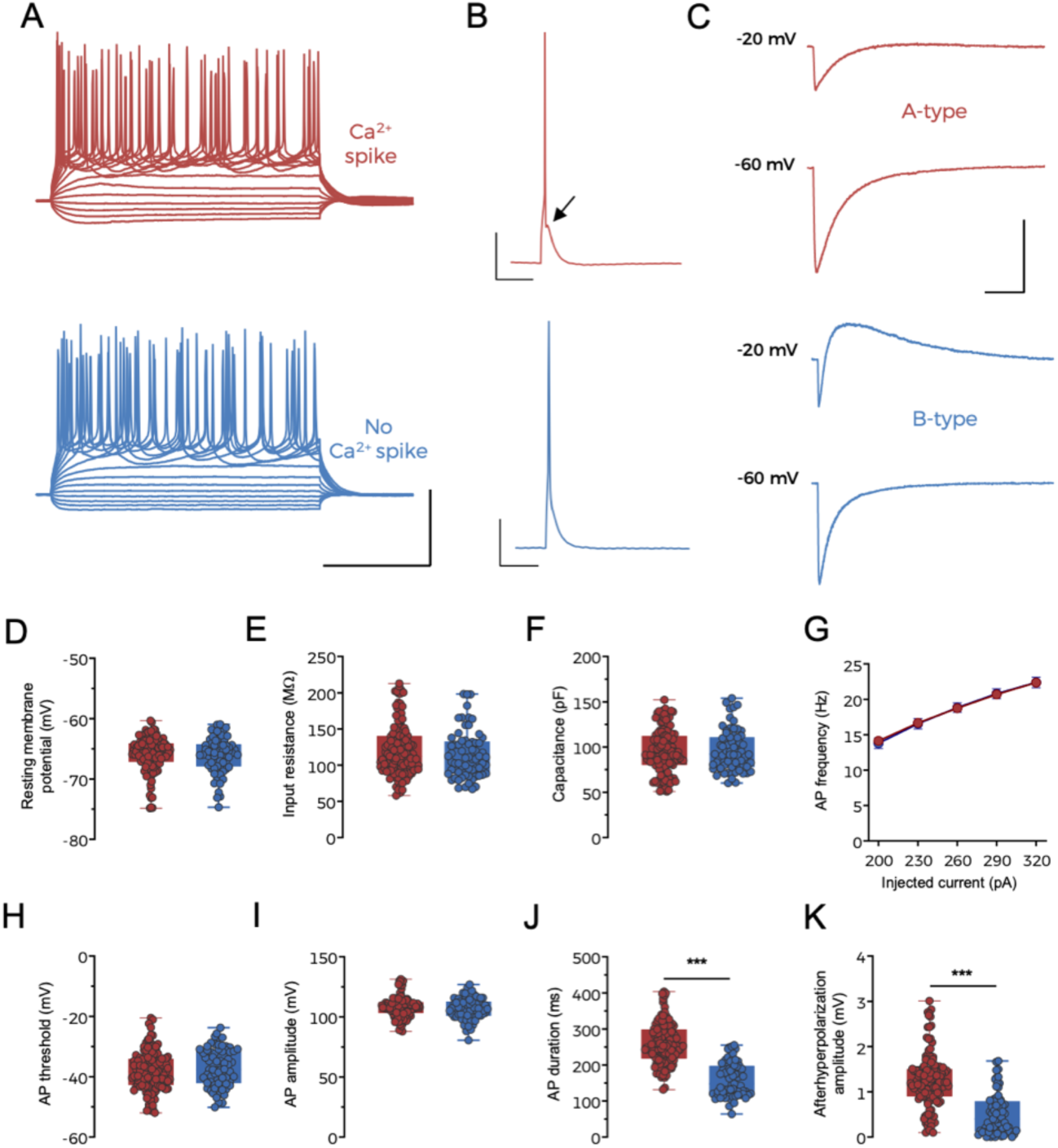
A subpopulation of regular-spiking BC-L5PNs exhibit NMDAR–BK functional coupling. **(A)** Representative current-clamp traces elicited with 500-ms depolarizing steps from -100 to 320 pA in A-type (red) and B-type (blue) BC-L5PNs. Scale bars represent 200 ms and 50 mV. **(B)** Single action potentials recorded from the same neurons shown in **(A)**. The arrow points to the Ca^2+^ spike apparent in A-type BC-L5PNs. Scale bars represent 100 ms and 20 mV. **(C)** NMDA-evoked currents recorded from the same neurons depicted in **(A)** and **(B)** at holding potentials of -20 and -60 mV with a Mg^2+^- free external solution containing 1 µM TTX and 10 µM glycine. Scale bars represent 5 s and 150 pA. **(D–K)** Values for resting membrane potential **(D)**, input resistance **(E)**, cell capacitance **(F)**, action potential frequency **(G)**, action potential threshold **(H)**, action potential amplitude **(I)**, action potential duration **(J)**, and afterhyperpolarization amplitude **(K)**. Data correspond to individual measurements (symbols) and median and 25^th^-75^th^ percentiles (boxes) from A-type (red, n = 130) and B-type (blue, n = 67) BC- L5PNs, except for **(G)**, where data points represent mean ± SEM. In **(J)** and **(K)**, ***p<0.001 (B-type vs. A-type). See also Figure S5 and Table S1.

### BK-dependent inhibition of NMDARs reduces postsynaptic response amplitude

Having demonstrated that BK channels in the basal dendrites of BC-L5PNs are activated after Ca^2+^ entry through proximal NMDARs, we predicted that this coupling would give rise to a negative feedback loop similar to that described for the coupling of VGCCs and BK channels in presynaptic terminals (Griguoli et al., 2016) or NMDARs and small-conductance Ca^2+^-activated K^+^ channels (SK) in postsynaptic terminals (Ngo-Anh et al., 2005). In such a scheme, NMDAR activation and subsequent entry of Ca^2+^ would open BK channels, which would repolarize the membrane due to outward flux of K^+^ and therefore reinstate voltage-dependent Mg^2+^ block of NMDARs, in turn truncating Ca^2+^ entry. If that were the case, the presence of postsynaptic NMDAR–BK coupling in B-type neurons should have a significant impact on synaptic transmission. Indeed, our finding that BK activation reduced NMDAR-mediated inward current in B-type neurons (Figure 1D) suggested the presence of such a negative feedback loop. To investigate the effect of NMDAR–BK coupling on synaptic transmission, we electrically stimulated the afferent inputs to BC-L5PN basal dendrites and recorded evoked postsynaptic potentials (PSPs) in physiological conditions (with Mg^2+^ in the external solution). Presynaptic stimulation of basal afferent inputs was performed at the limit between layers 5 and 6 to activate ascending thalamocortical fibers (Manns et al., 2004; Nunez et al., 2012), many of which are known to make direct contacts with BC-L5PNs (Agmon and Connors, 1992; Constantinople and Bruno, 2013; El-Boustani et al., 2020; Rodriguez-Moreno et al., 2020). The presence or absence of a Ca^2+^ spike in evoked action potentials allowed us to classify BC-L5PNs as either A-type or B-type. Stimulation intensity was then adjusted to obtain 3-5 mV PSPs in all recorded neurons (Figure 5A). Consistent with our proposed mechanism, BK channel block using 1 µM paxilline increased NMDAR availability in B-type neurons, thus increasing PSP amplitudes and slowing PSP kinetics (Figure 5B). Further addition of AP5 (100 µM) to the perfusate completely abolished the PSP amplitude increase and accelerated PSP kinetics (Figure 5B). This suggests that NMDAR-dependent activation of postsynaptic BK channels reduces the contribution of NMDARs to PSPs, thereby regulating synaptic transmission.

**Figure 5.**
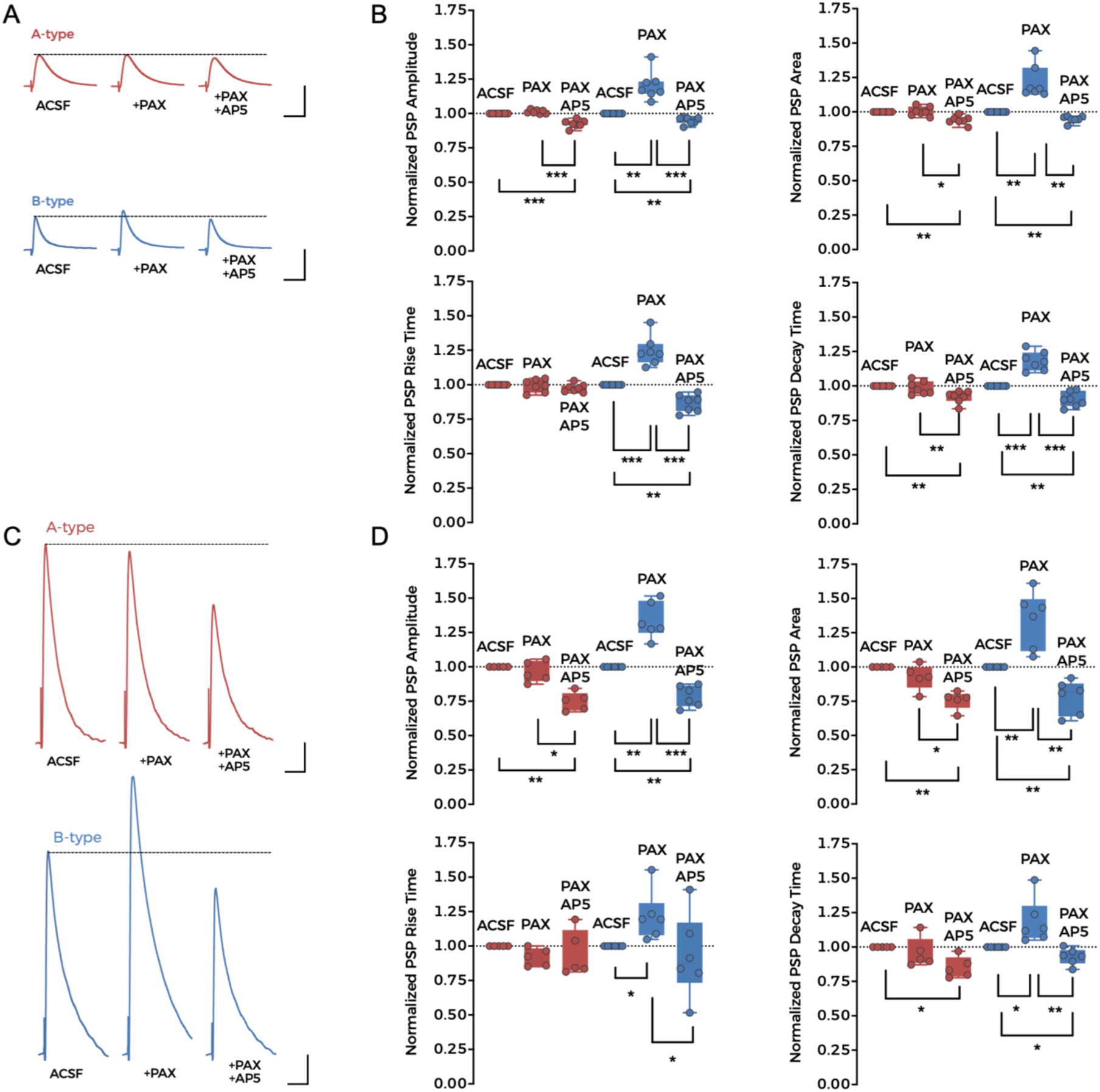
BK-dependent inhibition of NMDARs reduces postsynaptic response amplitude. **(A)** Representative synaptically-evoked postsynaptic potential (PSP) traces recorded from A-type (top, red) and B-type (bottom, blue) neurons in control conditions (ACSF, left), after the application of 1 µM paxilline (PAX, middle), and with 1 µM PAX plus 100 µM AP5 (PAX+AP5, right). Scale bars represent 50 ms and 3 mV. **(B)** Values for the normalized PSP amplitude (top left), area (top right), rise time (bottom left), and decay time (bottom right) in the experimental conditions in **(A)**. Data points represent individual measurements (n = 7 in both types of neuron) and boxes represent median and 25^th^-75^th^ percentile values. **(C)** Representative synaptically-evoked PSP traces recorded from A-type (top, red) and B-type (bottom, blue) neurons after increasing the electrical stimulation intensity applied to the afferent inputs. Conditions were the same as in **(A)** (ACSF, PAX, PAX+AP5). The pipette solution included 2 mM QX-314 to avoid action potential firing. Scale bars represent 50 ms and 3 mV. **(D)** Values for PSP amplitude (top left), area (top right), rise time (bottom left), and decay time (bottom right) for the experimental conditions shown in **(C)**. Data points represent individual measurements and boxes represent median and 25^th^-75^th^ percentile values. A-type: n = 5; B-type: n = 6. In **(B)** and **(D)**, *p<0.05, **p<0.01, and ***p<0.001. See also Table S1.

Coactivation of clustered neighboring basal inputs to BC-L5PNs initiates local dendritic NMDAR-dependent spikes that are characterized by large Ca^2+^ transients (Polsky et al., 2009; Schiller et al., 2000). We induced NMDA-dependent Ca^2+^ spikes by increasing the stimulation intensity applied to the afferent inputs to BC-L5PN basal dendrites while blocking action potentials by including the Na^+^ channel blocker QX-314 (2 mM) in the recording pipette. As expected, larger PSPs were observed in these experimental conditions (Figure 5C). Paxilline (1 µM) induced a further increase in PSP amplitude in B-type neurons, but had no effect on A-type neurons (Figures 5C and 5D). Further addition of AP5 (100 µM) reversed the PSP amplitude increase in B-type neurons to below control values and to a similar amplitude in both neuronal types (Figures 5C and 5D). These results demonstrate that BK channels in the basal dendrites of B-type neurons are able to abrogate NMDAR current, including under conditions where NMDAR spikes are taking place.

### NMDAR–BK coupling increases the threshold for induction of synaptic plasticity

The inhibitory effect of NMDAR**–**BK coupling on synaptic transmission in BC-L5PNs led us to wonder whether it may play a role in other physiological mechanisms, including forms of long-term synaptic plasticity involving NMDAR activation (Malenka and Nicoll, 1993; Markram et al., 1997). Spike-timing-dependent plasticity (STDP) is one such mechanism and relies on the precise coincidence of presynaptic and postsynaptic activity (Bi and Poo, 1998; Markram et al., 1997). The timing and order of presynaptic and postsynaptic action potentials determine the direction of the change in synaptic strength: a presynaptic action potential followed by a postsynaptic action potential within a window of tens of ms results in long-term potentiation (LTP), whereas the reverse order within a similar timeframe results in long-term depression (LTD) (Bi and Poo, 1998; Feldman, 2000; Markram et al., 1997). Both mechanisms are dependent on NMDAR activation (Nevian and Sakmann, 2006) but only spike-timing-dependent LTP (t-LTP) depends on postsynaptic NMDAR activation and the consequential rise in dendritic spine Ca^2+^ concentration (Nevian and Sakmann, 2006) (Figure S3). Because NMDAR activity is blunted by BK activation in the basal dendrites of B-type BC-L5PNs, we hypothesized that t-LTP would be less prominent in these neurons compared to A-type BC-L5PNs, or possibly absent. We therefore studied the effects of pairing pre- and postsynaptic action potentials in A-type and B-type BC- L5PNs.

We measured PSPs evoked by electrical stimulation of basal afferent inputs to BC- L5PNs before and after pre–post pairings at 0.20 Hz (Figure 6A). A low number of pre– post associations (30 pairings) induced t-LTP in A-type but not B-type neurons (Figure 6B, left). This is consistent with a reduction in the amount of Ca^2+^ entering the basal dendrites of B-type neurons due to NMDAR-dependent activation of BK channels. The difference between the two populations was abolished when 1 µM paxilline was included in the recording pipette (Figure 6B, right), suggesting that the release of NMDARs from BK-induced negative feedback favors t-LTP by reducing the threshold for its induction. The degree of t-LTP in both neuronal types in the presence of paxilline was significantly greater than that elicited in A-type neurons in control conditions (Figure 6B, right versus left), consistent with the key role of BK channels in the regulation of neuronal excitability (Latorre et al., 2017). Interestingly, these data imply that BK channels are functionally expressed in both A-type and B-type BC-L5PNs, which we confirmed by recording BK currents in both types of BC-L5PN (Figure S4). As our data indicated that NMDAR–BK coupling is restricted to the basal dendrites of BC-L5PNs, we asked whether the difference in synaptic plasticity threshold between A-type and B-type neurons was limited to basal dendrites, or could also occur in apical dendrites. We electrically stimulated the afferent inputs to the apical dendrites of BC- L5PNs (Figure 6C) and carried out 30 pre–post pairings at 0.20 Hz. This failed to induce synaptic plasticity in any of the neurons tested (Figure 6D, left), but when the number of pre–post pairings was increased to 90, t-LTP was induced in both types of neuron (Figure 6D, right). The potentiation was of a similar amplitude in both A- and B-type neurons, but significantly lower than that induced in basal dendrites by 30 pairings (compare Figure 6D, right with Figure 6B, left). These results confirm that functional NMDAR–BK coupling occurs exclusively in the basal dendrites of B-type BC-L5PNs, where it increases the threshold for synaptic plasticity and therefore modulates neuronal circuits involving these dendrites.

**Figure 6.**
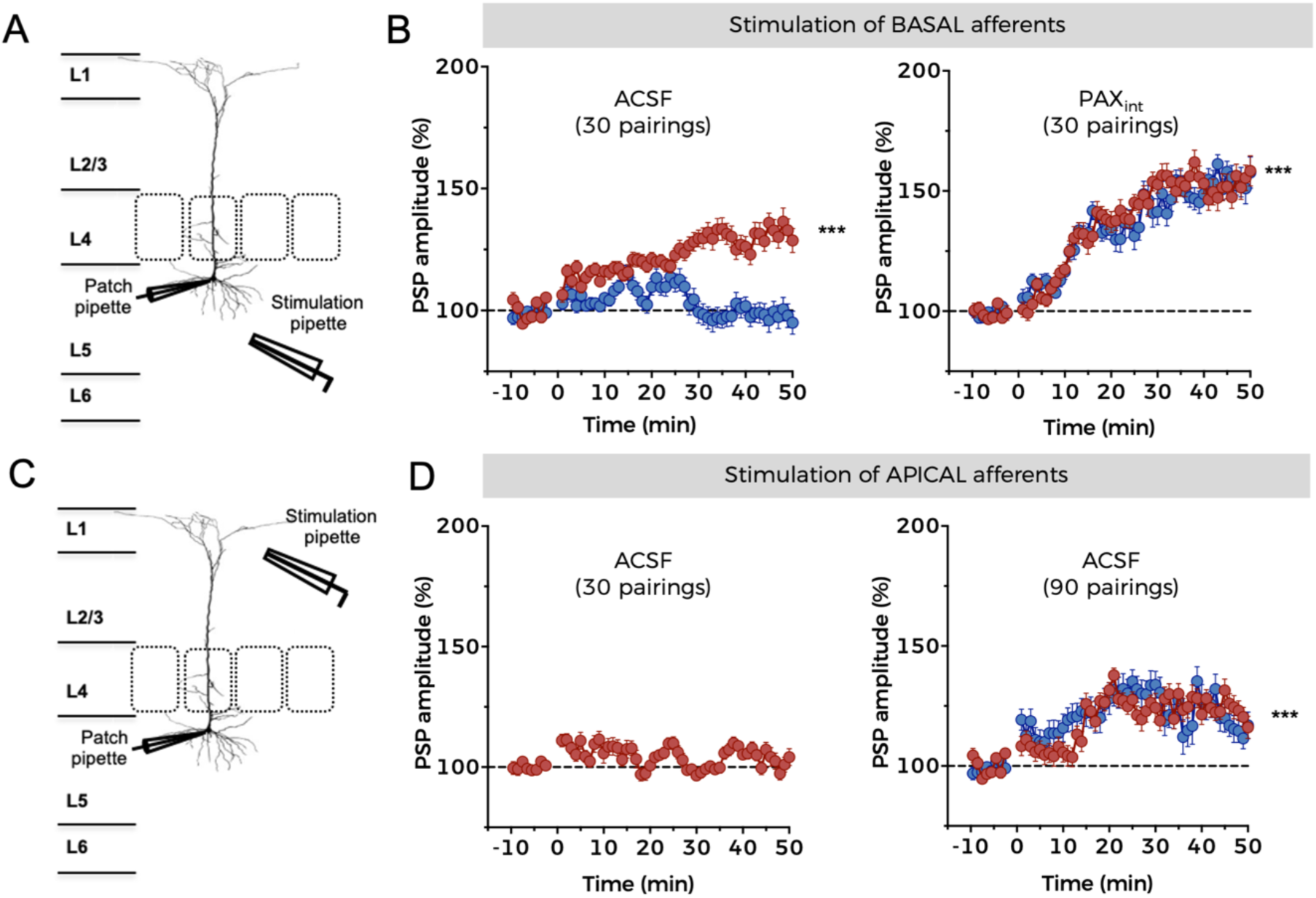
NMDAR–BK coupling increases the threshold for induction of synaptic plasticity. **(A)** Schematic representation of the experimental design used to induce t-LTP via stimulation of basal afferent inputs to BC-L5PNs. **(B)** Development of t-LTP over time in A-type (red) and B-type (blue) neurons in control conditions (ACSF, left) and in the presence of 500 nM intracellular paxilline (PAX, right). Data points represent mean ± SEM. A-type (ACSF): n = 6; B-type (ACSF): n = 4; A-type (PAX): n = 6; B-type (PAX): n = 4. ***p<0.001 (t-LTP vs. basal conditions). **(C)** Schematic representation of the experimental design used to induce t-LTP via stimulation of apical afferent inputs to BC-L5PNs. **(D)** Development of t-LTP over time in A-type (red) and B-type (blue) neurons in control conditions (ACSF) using 30 (left) or 90 (right) STDP pairings. Data points represent mean ± SEM. A-type (30 pairings): n = 3; A-type (90 pairings): n = 6; B-type (90 pairings): n = 3. See also Figure S3, S4, and Table S1.

### A high number and frequency of pre–post pairings relieves BK-dependent NMDAR inhibition

Our results indicated that BK reduces, but does not completely abolish, the influx of ions through NMDARs, and therefore increases the threshold for the induction of synaptic plasticity (Figure 6B). We reasoned that by tuning the experimental conditions to induce greater NMDAR activation, the concentration of Ca^2+^ in postsynaptic terminals would eventually reach sufficient levels to induce plasticity in B-type BC-L5PNs. In A-type neurons, increasing the number of pre–post pairings to 50 or 90 increased the extent of t-LTP, as previously described in other brain areas including the hippocampus (Fernandez de Sevilla and Buno, 2010; Wittenberg and Wang, 2006; Zhang et al., 2009). Importantly, we were also able to induce t-LTP in B-type neurons by increasing the number of pre–post associations (Figure 7A), confirming that sufficient NMDAR activation can overcome the higher threshold for plasticity in these neurons. However, the extent of PSP potentiation in B-type neurons was significantly less than in A-type neurons for a given number of pairings (Figure 7B). These results therefore demonstrate that NMDAR–BK coupling regulates the threshold to induce synaptic plasticity in the basal dendrites of B-type BC-L5PNs.

**Figure 7.**
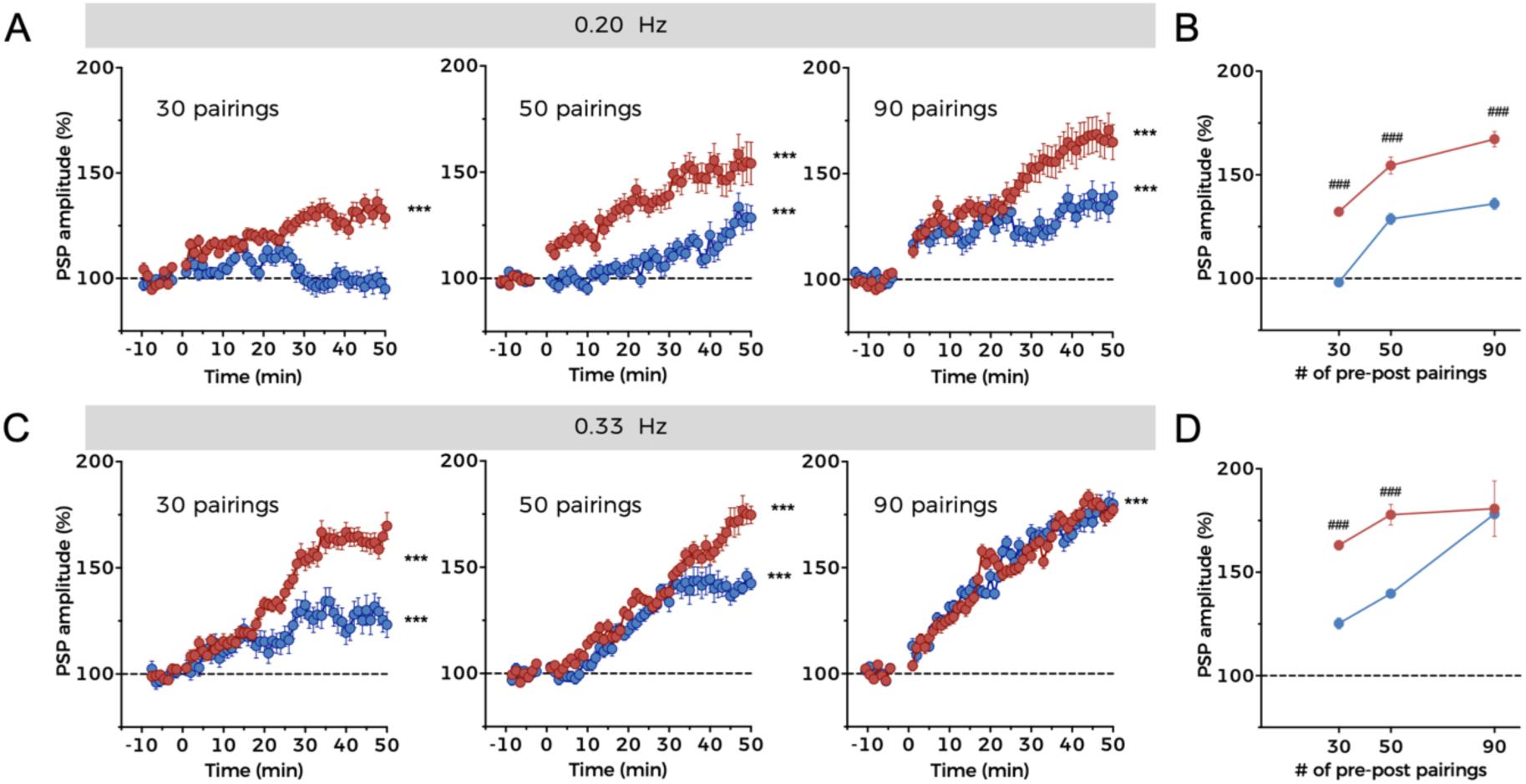
A high number and frequency of pre–post pairings relieves BK- dependent NMDAR inhibition. **(A)** Time course of t-LTP development over time in A-type (red) and B-type (blue) neurons using 30 (left), 50 (middle), or 90 (right) pairing STDP protocols at frequency of 0.20 Hz using the experimental design represented in Figure 6A. Data points represent mean ± SEM. A-type (30 pairings): n = 6, B-type (30 pairings): n = 4; A-type (50 pairings): n = 6; B-type (50 pairings): n = 4; A-type (90 pairings): n = 6; B-type (90 pairings): n = 5. Data in the left panel (30 pairings) are the same as in Figure 6B. **(B)** Summary of t-LTP extent for A-type (red) and B-type (blue) neurons under the experimental conditions depicted in **(A)**. **(C)** Time course of t-LTP development over time in A-type (red) and B-type (blue) neurons using 30 (left), 50 (middle), or 90 (right) pairing STDP protocols at a frequency of 0.33 Hz. Data points represent mean ± SEM. A-type (30 pairings): n = 6, B-type (30 pairings): n = 4; A-type (50 pairings): n =7; B-type (50 pairings): n = 5; A-type (90 pairings): n = 5; B-type (90 pairings): n = 6. **(D)** Summary of t-LTP extent for A-type (red) and B-type (blue) neurons under the experimental conditions depicted in **(C)**. In **(A)** and **(C)**, ***p<0.001 (t-LTP vs. basal conditions). In **(B)** and **(D)**, ^###^p<0.001 (B-type vs. A-type). See also Table S1.

The magnitude of t-LTP can be also regulated by changing the frequency or modifying the time window of pairings (Caporale and Dan, 2008; Nevian and Sakmann, 2006), although the latter results in considerable STDP variability depending on the synapse studied (Caporale and Dan, 2008). We hypothesized that by increasing the frequency of pairings, a condition would be reached where the inhibitory effect of BK on NMDARs completely disappears. For this purpose, we applied 30, 50, and 90 pairings at a frequency of 0.33 Hz (pre–post pairings delivered every 3 s), and found that the degree of t-LTP in each condition was greater than that at 0.20 Hz for both types of neuron (compare Figure 7C with Figure 7A). Furthermore, there was no difference in the extent of PSP potentiation between A-type and B-type neurons following the delivery of 90 pairings (Figure 7D). Thus, the inhibitory effect of BK on NMDARs can be abolished with high rates of pre- and postsynaptic coincident activity. Moreover, although t-LTP appeared to saturate following 50 pairings in A-type neurons subjected to 0.33 Hz stimulation, B-type BC-L5PNs remained able to discriminate between the number of pairings in the protocol.

## DISCUSSION

We have described the functional association of NMDARs and BK channels, and the role of this coupling in synaptic function, in the basal dendrites of a population of regular-spiking BC-L5PNs. NMDAR activation and subsequent Ca^2+^ entry promotes BK opening, which repolarizes the cell membrane and halts NMDAR activity. Both GluN2A- and GluN2B-containing NMDARs can activate BK channels, although our data suggest that GluN2B-containing NMDARs are more efficient. Functional coupling of NMDARs and BK channels in the basal dendrites of this specific BC-L5PN population modulates synaptic transmission and produces an increase in the threshold for induction of synaptic plasticity, indicating that NMDAR–BK association critically influences how specific neuronal types integrate afferent synaptic inputs. In fact, NMDAR–BK functional coupling bestows B-type BC-L5PNs with the ability to work as high-pass filters of incoming inputs, depending on the number and frequency of afferent stimuli.

### The molecular basis of NMDAR–BK functional coupling

In the CNS, BK channels can couple to different Ca^2+^-conducting channels, including VGCCs (Marrion and Tavalin, 1998), RyRs (Chavis et al., 1998), and InsP3Rs (Zhao et al., 2010). Such Ca^2+^ sources provide the Ca^2+^ needed for BK activation, but membrane depolarization is generally provided by a coincident action potential (Griguoli et al., 2016). Interestingly, NMDAR activation can provide both the membrane depolarization and the Ca^2+^ entry required for BK activation (Paoletti et al., 2013), particularly in restricted compartments such as the dendritic spine, where Ca^2+^ concentrations reach micromolar levels after NMDAR activation (Higley and Sabatini, 2012; Sabatini et al., 2002). In contrast to previous studies using Mg^2+^-containing ACSF (Isaacson and Murphy, 2001; Zhang et al., 2018), we initially observed functional association of NMDARs and BK channels in the absence of Mg^2+^. These data revealed that NMDAR-dependent BK activation occurs at potentials positive to - 40 mV, within a range of potentials at which the NMDAR-dependent current is maximal (Mayer et al., 1984; Nowak et al., 1984). This was subsequently corroborated in synaptic transmission experiments performed in the presence of Mg^2+^, in which inhibition of NMDARs by BK channels became evident.

Consistent with previous studies (Isaacson and Murphy, 2001; Zhang et al., 2018), our results showed that NMDAR-mediated increases in Ca^2+^ concentration are required in the immediate vicinity of BK channels in order to influence their activation. Thus, the coupling mechanism relies on the close proximity of NMDARs and BK channels in the plasma membrane. Our experiments using Ca^2+^ chelators allowed us to estimate that the two proteins must be situated within 15-60 nm of each other for functional coupling to occur. However, this estimate must be taken with caution, as the exact concentration of chelators that reach dendrites is not known. In fact, our experiments using PLAs suggest that the maximum distance between the two channels may be even shorter (below 40 nm). Co-immunoprecipitation and biochemical approaches have indicated direct association of NMDARs and BK channels in the soma (Zhang et al., 2018). Our results demonstrate strong functional coupling between NMDARs and BK channels in basal dendrites of BC-L5PNs, regardless of whether the channels physically interact with each other.

### NMDAR–BK functional coupling regulates long-term synaptic plasticity

Our work provides the first evidence for a functional role of NMDAR–BK coupling in neuronal dendrites. This result differs from previous studies in which functional coupling between NMDARs and BK channels has been observed in the soma of granule cells from the olfactory bulb (Isaacson and Murphy, 2001) and dentate gyrus (Zhang et al., 2018). Some evidence also points to NMDAR–BK interactions in hippocampal CA1 pyramidal neuron somata (Zorumski et al., 1989) but not dendrites ((Ngo-Anh et al., 2005) and Figure S2). At the soma, NMDAR-dependent activation of BK channels would likely constitute a mechanism that regulates action potential shape, and controls neuronal excitability, independently of dendritic input.

In dendrites, NMDAR–BK coupling would generate a negative feedback mechanism that could have dramatic effects on forms of synaptic plasticity that involve NMDARs and Ca^2+^ entry. Here we have demonstrated that B-type BC-L5PNs exhibiting NMDAR–BK functional coupling have a higher threshold for the induction of long-term synaptic plasticity. This phenomenon of selective plasticity attenuation is restricted to the basal dendrites of these neurons, as the effect was not observed when afferent inputs to apical dendrites were stimulated. Similar basal versus apical polarity differences that affect synaptic input integration have been described in other brain areas, including the hippocampus (Cornford et al., 2019).

Interestingly, a recent study suggests that SK-mediated inhibition of NMDARs is a general mechanism that regulates synaptic plasticity associated with BC-L5PN to BC- L5PN communication (Jones et al., 2017). This backward regulatory mechanism affects intralayer communication between regular-spiking BC-L5PNs and depends on back-propagating action potentials rather than NMDA activation. Although we cannot exclude a contribution from this process, there are three lines of evidence to suggest that the majority of effects we describe are due to NMDAR–BK coupling: (i) NMDAR activation resulted only in BK channel-mediated current; (ii) BK-specific inhibition completely abolished the t-LTP differences between A-type and B-type BC-L5PNs; and (iii) BK-mediated inhibition of NMDARs remained when basal afferent inputs were electrically stimulated and was independent of back-propagating action potentials. Therefore, although two different mechanisms for inhibition of NMDARs may be present in the basal dendrites of BC-L5PNs, our data strongly suggest that NMDAR– BK modulation of synaptic transmission and long-term synaptic plasticity is a forward regulatory mechanism involving thalamocortical projections to a restricted population of BC-L5PNs in conditions where prior activation of postsynaptic NMDARs is mandatory and action potentials aren’t required.

We used a low frequency STDP protocol to induce LTP in our study. Under these experimental conditions, it has been proposed that GluN1/GluN2B NMDARs make a larger contribution to the total charge transfer than GluN1/GluN2A NMDARs (Erreger et al., 2005), as expected from the slower deactivation rates of GluN2B-containing NMDARs (Vicini et al., 1998). Taking this into account, GluN2B-containing NMDARs should conduct more Ca^2+^ than GluN1/GluN2A channels in our experimental conditions, activating BK more efficiently. That being the case, Ca^2+^ entry through GluN2B-containing NMDARs would be the main contributor to BK activation and thus the inhibitory mechanism underlying modulation of synaptic transmission and LTP. This notion corresponds with our heterologous expression experiments using physiological concentrations of extracellular Na^+^, in which GluN1/GluN2B NMDARs produced a larger leftward shift in the BK activation curve than GluN1/GluN2A. It also correlates with our observations in basal dendrites of BC-L5PNs, where specific blockade of GluN2B-containing NMDARs produced a larger reduction in the NMDA- evoked outward current. In summary, we have demonstrated that GluN2A- and GluN2B-containing NMDARs are able to activate BK channels in both heterologous expression systems and the basal dendrites of BC-L5PNs. Whether B-type neurons express a combination of GluN1/GluN2A and GluN1/GluN2B heteromers, or GluN1/GluN2A/GluN2B triheteromers (Paoletti et al., 2013), requires further investigation.

### Regular-spiking BC-L5PNs act as high-pass filters

An interesting question arises from the observation that functional NMDAR–BK association is exclusive to the basal dendrites of a subpopulation of regular-spiking BC-L5PNs: is it associated with the specific expression of particular GluN2 subunits? Although the role of GluN2C and GluN2D subunits was not investigated, a differential distribution of these, and other, subunits between A-type and B-type neurons would be reflected in the macroscopic NMDAR conductance, which is similar in both A-type and B-type BC-L5PNs. Therefore, we believe that both populations of BC-L5PNs exhibit a similar distribution of GluN2 subunits. Why, then, do NMDAR–BK associations occur only in the basal dendrites of B-type neurons? As both cell types express BK channels and NMDARs, a plausible explanation is that a mechanism exists to target specific channels to dendritic compartments. This could be achieved by engaging scaffolding proteins, such as the receptor for activated C kinase 1 (RACK1) and caveolin-1, which are known to bind both the GluN2B NMDAR subunit (Yaka et al., 2002; Yang et al., 2015) and BK channels (Isacson et al., 2007; Wang et al., 2005). In addition, we observed a larger BK current in B-type than in A-type neurons, suggesting a higher abundance of BK channels in their membrane. Therefore, it is tempting to speculate that the formation of complexes and their targeting to the basal dendrites of B-type neurons depends on the overall abundance of BK channels and thus their availability to couple to NMDARs.

In this study, we uncover two populations of regular-spiking BC-L5PNs that are distinguished by the absence (A-type, ∼64%) or presence (B-type, ∼36%) of NMDAR– BK functional coupling in basal dendrites. Interestingly, this distribution resembles the electrophysiological characteristics and neuronal ratio of two previously described populations of mouse and rat BC-L5PNs, which were classified according to the presence or absence of a Ca^2+^ spike (Agmon and Connors, 1992; Chagnac-Amitai and Connors, 1989; Maglio et al., 2018). Functional NMDAR–BK coupling is exclusive to the basal dendrites of B-type neurons (Figure 8), which exhibit a higher threshold for the induction of LTP due to BK-dependent inhibition of NMDARs. A-type neurons lack this molecular brake, and therefore reach saturation at a lower stimulation frequency, independent of the number of synaptic inputs. This leads us to propose that BK- dependent inhibition of NMDARs endows B-type neurons with a calibration mechanism that allows them to decode the number and frequency of afferent synaptic inputs using selective synaptic plasticity attenuation. As a result of this discrimination capability, we hypothesize that the basal dendrites of B-type BC-L5PN function as high-pass filters of thalamic afferent inputs, displaying the same output as A-type neurons when a strong stimulus or series of stimuli reach the dendrites (such as during high levels of pre- and postsynaptic coincident activity), but attenuating signals below a specific stimulation threshold. This cutoff would be mainly determined by the number of BK channels that are available to functionally couple to NMDARs in the basal dendrites of B-type neurons: the larger the number of available BK channels, the higher the threshold for induction of synaptic plasticity. This mechanism would provide B-type BC- L5PNs with a dynamic range of output responses for the same afferent input stimuli, thus increasing the computational power of the somatosensory cortex.

**Figure 8.**
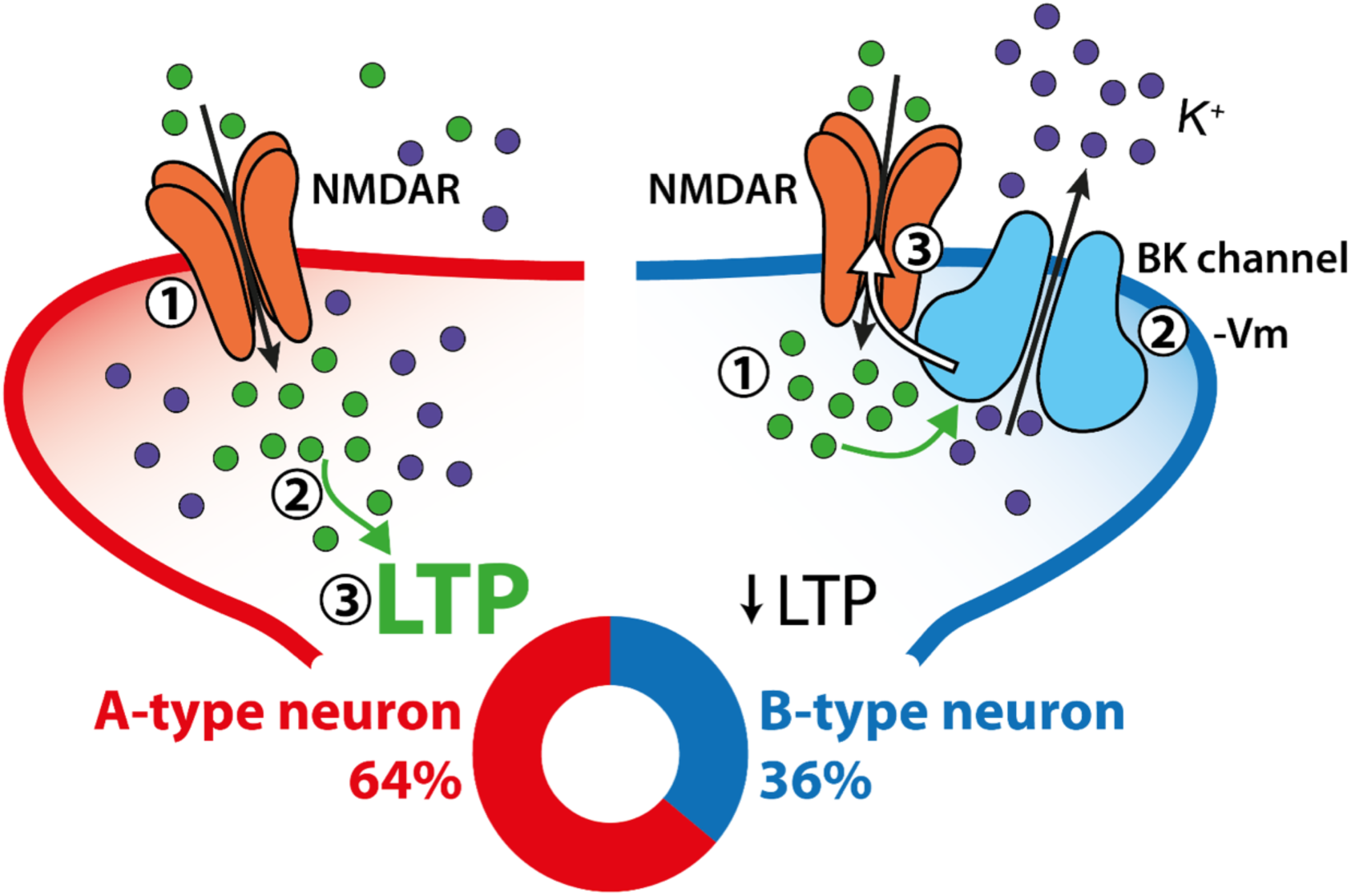
NMDAR–BK coupling controls dendrite-specific synaptic plasticity. Two populations of regular-spiking BC-L5PNs can be distinguished by the absence (A-type, left, red) or presence (B-type, right, blue) of NMDAR-BK functional association in basal dendrites. In B-type neurons (∼36%), NMDAR–BK coupling provides a negative feedback mechanism whereby the entry of Ca^2+^ associated with NMDAR activation **(1)** opens neighbouring BK channels **(2)** that allow outward flow of K^+^. The resultant membrane hyperpolarization **(-Vm)** reinstates voltage-dependent Mg^2+^ block of NMDARs **(3)**, truncating Ca^2+^ entry and increasing the threshold for long-term synaptic plasticity **(↓LTP**). We show in this study that B-type BC-L5PNs exhibit a higher threshold for induction of t-LTP. On the other hand, A-type neurons (∼64%) lacking the NMDAR–BK molecular break undergo long-term potentiation (**3**) associated with Ca^2+^ entry **(2)** via NMDARs **(1).** Our data reveal that A-type neurons reach saturation at a lower stimulation frequency than B-type, independent of the number of synaptic inputs.

In summary, we have demonstrated that functional coupling of NMDARs and BK channels in the basal dendrites of a specific set of BC-L5PNs modulates synaptic transmission and synaptic plasticity in thalamocortical circuits. This finding unmasks the critical influence that functional association of ion channels can have on the integration of afferent synaptic inputs by neurons.

## Supporting information

Supplementary text and Figures Gomez et al

## ACKNOWLEDGMENTS

Authors thank Drs. A. J. Plested, W. Buño, J. S. Diamond and A. Pérez-Álvarez for their critical reading of the manuscript. This work has been funded by the European Research Council, under Horizon 2020 Research and Innovation Programme Grant (ERC-CoG-2014 648936) and Spanish Ministerio de Ciencia, Innovación y Universidades (RTI2018-098768-B-I00) to TG. A.J.G.-H. received a Predoctoral Fellowship from the Spanish Ministerio de Educación, Cultura y Deporte (FPU15/02528).

## AUTHOR CONTRIBUTIONS

Conceptualization, R.G., L.E.M., and T.G.; methodology, R.G. and L.E.M.; formal analysis, R.G., L.E.M., A.J.G-H., B.R-P., and D.B-M.; investigation, R.G., L.E.M., A.J.G-H., B.R-P., and D.B-M.; writing original draft, R.G. and T.G.; draft review & editing, R.G. and T.G.; funding acquisition, T.G. All authors read, discussed, and approved the manuscript.

## MATERIAL and METHODS

### KEY RESOURCES TABLE

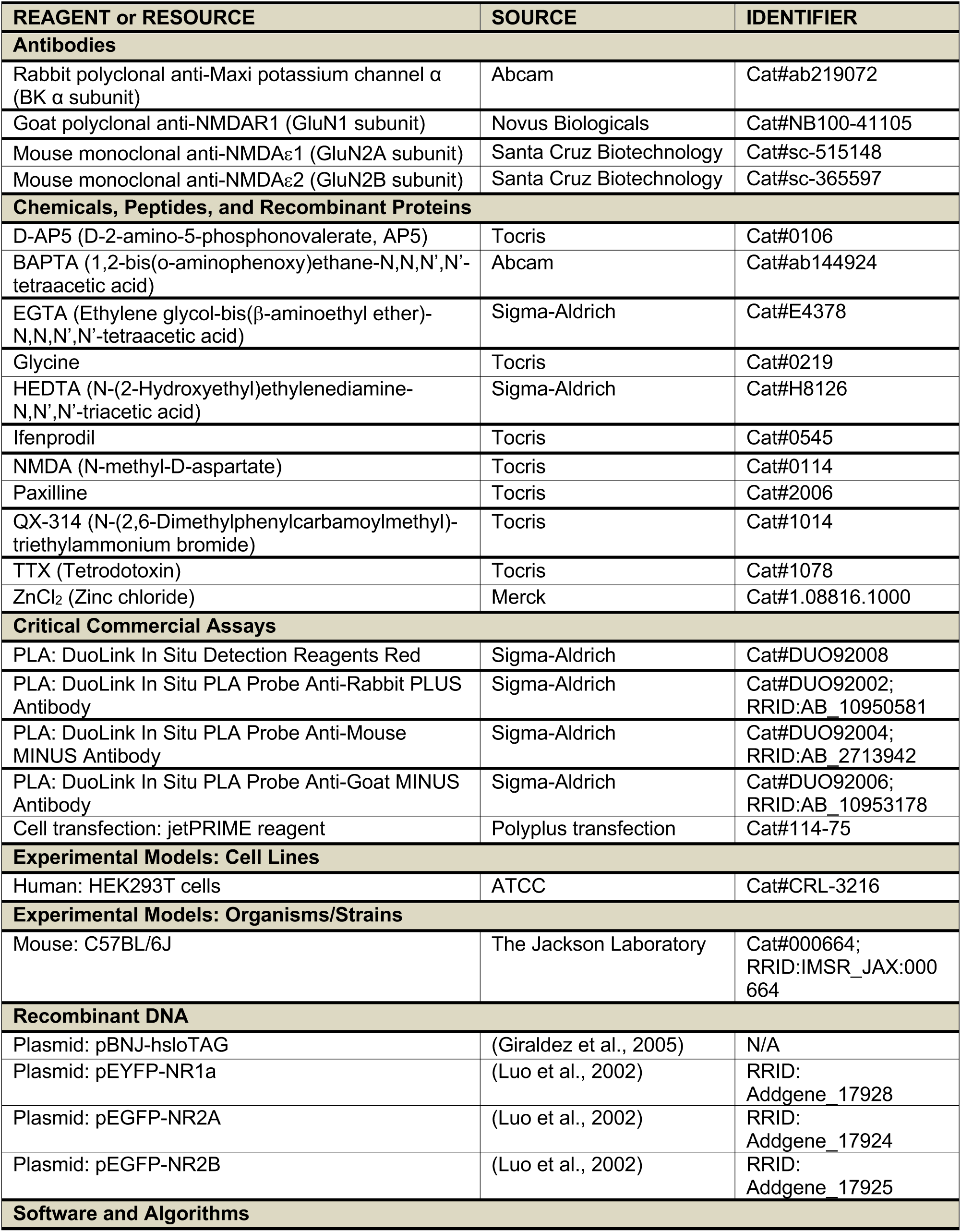
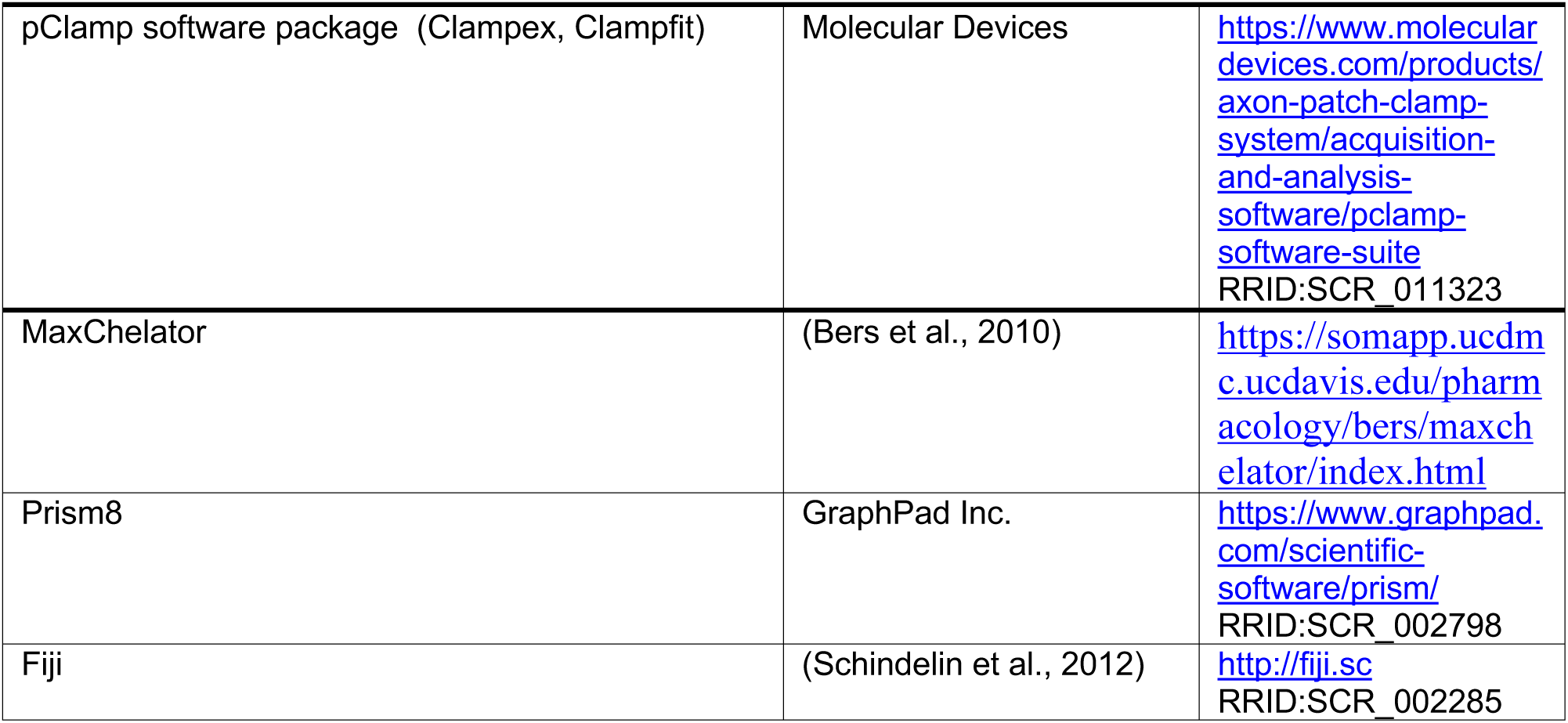

### LEAD CONTACT AND MATERIALS AVAILABILITY

Further information and requests for reagents and resources should be directed to the lead contact author, Teresa Giraldez.

### EXPERIMENTAL MODEL

All the procedures for handling and sacrificing animals followed the Spanish (Real Decreto 53/2013) and European Commission (Directive 2010/63/EU) guidelines for the care and use of laboratory animals and were approved by the Ethics Committee (CEIBA) of the Universidad de La Laguna. C57BL/6J mice (The Jackson Laboratory #000664) were housed in standard laboratory cages with *ad libitum* access to water and food in temperature- and humidity-controlled rooms under a 12:12 h light-dark cycle at the Universidad de La Laguna animal facilities (registry number ES380230024514).

### METHOD DETAILS

#### Acute mice brain slices

C57BL/6J adult male mice (postnatal 28-36) were slightly anesthetized with isoflurane, decapitated, and the brain quickly removed and immersed in an ice-cold high-sucrose “cutting solution” (composition in mM: 189 sucrose, 10 glucose, 26 NaHCO_3_, 3 KCl, 5 MgSO_4_, 0.1 CaCl_2_, and 1.25 NaH_2_PO_4_), continuously bubbled with carbogen (95% oxygen/5% carbon dioxide). Coronal brain slices of 350 µm of thickness were obtained using a Microm HM650V vibratome (Thermo Scientific). These slices were maintained during 1-1.5 h in artificial cerebrospinal fluid (ACSF; composition in mM: 124 NaCl, 2.69 KCl, 1.25 KH_2_PO_4_, 2 MgSO_4_, 26 NaHCO_3_, 10 glucose, 2 CaCl_2_, and 0.4 ascorbic acid; pH 7.35) bubbled with carbogen at room temperature (22-24°C) for recovery. After recovery, brain slices were transferred to a recording chamber mounted onto an upright Olympus BX51WI microscope (Olympus) equipped with 10x/40x water immersion objectives, where they were continuously perfused with carbogen-bubbled ACSF.

#### Electrophysiology in brain slices

Electrophysiological recordings in brain slices were performed using the whole-cell configuration of the patch-clamp technique (Hamill et al., 1981). Patch pipettes were made from 1.50 OD/0.86 ID borosilicate glass capillaries (GC150F-10; Harvard Apparatus) using a P-97 micropipette puller (Sutter Instruments) and had resistances of 5-8 MΩ when filled with the internal solution (composition in mM: 135 KMeSO_4_, 10 KCl, 10 HEPES, 5 NaCl, 2.5 ATP-Mg, and 0.3 GTP-Na; pH 7.3). Recordings were obtained with a MultiClamp 700A amplifier (Molecular Devices). Series resistance was compensated (≍70%) and recordings were rejected when the access resistance varied >20% during the experiment. Data were acquired at 10 kHz and lowpass-filtered at 4 kHz. Signals were fed to a Pentium-based PC through an Axon Digidata 1550B interface board (Molecular Devices). The pClamp software (Molecular Devices) was used for stimulus generation, data display, acquisition, storage, and analysis. All experiments were performed at room temperature (22-24°C).

Pyramidal neurons from layer 5 of the somatosensory 1 barrel field area (S1BF, namely “barrel cortex”) were visualized by infrared microscopy and differential interference contrast (IR-DIC) using an ORCA-Flash4.0LT digital camera (Hamamatsu). Barrel cortex layer 5 pyramidal neurons (BC-L5PN) were identified by their location (just below layer 4), shape (characteristic somatic morphology), and electrophysiological properties (only regular-spiking neurons were used in the study; intrinsically-bursting and fast-spiking neurons were discarded), suggesting that we predominantly recorded from slender-tufted neurons located in layer 5a, as described previously (Manns et al., 2004; Nunez et al., 2012).

##### Recording of N-methyl-D-aspartate-evoked currents

N-methyl-D-aspartate (NMDA; 200 μM) was locally delivered (“puff” application) at different dendritic locations of BC-L5PN through a glass pipette using a PMI-100 Pressure MicroInjector (Dagan Corporation) at 1 bar during 50-200 ms (Figure 1A and Figure Supp. 1A). Puff pipettes were made from GC150F-10 borosilicate glass capillaries as patch pipettes. Responses evoked by NMDA application at different holding potentials (from -60 to 0 mV) were recorded from BC-L5PN using the voltage-clamp mode in the whole-cell configuration of the patch-clamp technique in a Mg^2+^- free ACSF, where an equimolar substitution of Mg^2+^ with Ca^2+^ was made to maintain total extracellular divalent cation concentration. The Mg^2+^-free ACSF was supplemented with tetrodotoxin (TTX; 1 µM) to avoid neuronal action potential-dependent communication and with the NMDAR coagonist glycine (10 µM) to ensure NMDAR activation. The duration of the NMDA puff application was adjusted to obtain inward currents of 200-250 pA of peak amplitude at -60 mV. NMDA local application was performed at different BC-L5PN locations: basal dendrites (Figure 1), oblique dendrites/initial segment of the apical dendrite (Figure Supp. 1), and apical dendrites (not shown). We were unable to record NMDA-evoked currents when NMDA was applied at apical dendrites of BC-L5PN under our experimental conditions, which may suggest that we predominantly recorded from slender-tufted neurons. In some experiments, responses evoked by NMDA application at different holding potentials (from -70 to -10 mV) were recorded from hippocampal CA1 pyramidal neurons using the same experimental conditions described above (Figure Supp. 4).

NMDA-evoked inward currents (I_Inward_, namely I_NMDAR_) were measured as the maximum peak amplitude, whereas NMDA-evoked inward and outward total ionic charges (Q_Inward_ and Q_Outward_, respectively; namely Q_NMDAR_ and Q_BK_) were calculated as the area under the recorded trace and the zero current level. The I_NMDAR_, Q_NMDAR_, and Q_Outward_/Q_Inward_ ratio were calculated and represented as a function of the holding potential (Figures 1C and 1D). BC-L5PN were classified according to the absence (A-type neurons) or presence (B-type neurons) of an NMDA-evoked outward current.

Pharmacological characterization of inward and outward currents was performed in B-type neurons at a holding potential of -20 mV (Figures 1E and 1F). All drugs were added to the bath solution. In some experiments, BAPTA (15 or 1 mM) and EGTA (15 mM) Ca^2+^ chelators were added to the recording pipette (Figure 2), where an equimolar substitution of KMeSO_4_ with BAPTA-K or EGTA-K was made to maintain the total internal K^+^ concentration.

##### Action potential and postsynaptic potential recordings

BC-L5PN were recorded using the current-clamp mode in the whole-cell configuration of the patch-clamp technique in normal ACSF (2 MgSO_4_ and 2 CaCl_2_). Patch pipettes and internal solution were the same as described previously for voltage-clamp recordings.

Current-voltage (I-V) relationships were elicited with 500-ms depolarizing steps from - 100 to 320 pA (Figure 4A). Resting membrane potential, action potential firing, and cell input resistance were calculated from these recordings (Figure 4D to 4G).

Action potentials were obtained either by presynaptic electrical stimulation of basal afferent inputs or by a brief current injection through the recording pipette (5 ms, 200- 400 pA) since no differences between both methods were found (see Figure Supp. 5A for a comparison). Action potential characteristics were calculated from these recordings as depicted in Figures Supp. 5C and 5D. According to the absence or presence of a Ca^2+^ spike following the action potential, two different populations of BC- L5PN were distinguished, as previously described (Agmon and Connors, 1992; Chagnac-Amitai and Connors, 1989; Maglio et al., 2018). These populations inversely correlated with those described after NMDA-evoked current recordings, that is, A-type neurons exhibited Ca^2+^ spikes but no NMDAR-evoked outward currents, whereas B-type neurons exhibited NMDAR-evoked outward currents but no Ca^2+^ spikes (Figure 4B and 4C).

Postsynaptic potentials (PSP) were obtained by presynaptic electrical stimulation of basal afferent inputs in the limit between layers 5 and 6 since it is widely described that it activates an important contribution of ascending thalamocortical fibers (Manns et al., 2004; Nunez et al., 2012), many of which are known to directly connect with BC-L5PN (Agmon and Connors, 1992; Constantinople and Bruno, 2013; El-Boustani et al., 2020; Rodriguez-Moreno et al., 2020). Stimulation pipettes were made from 1.50 OD/1.02 ID thick septum theta borosilicate glass capillaries (TST150-6; World Precision Instruments) using a P-97 micropipette puller and filled with ACSF. Single pulses (100 µs) were delivered at 0.20 Hz by a Master-8 pulse Stimulator (A.M.P.I.) through an ISU-165 isolation unit (Cibertec). Stimulus intensity was adjusted to obtain 3-5 mV PSP responses and was unchanged for the entire experiment (Figure 5A). In some experiments (Figure 5C), stimulus intensity was adjusted to evoke NMDAR-dependent Ca^2+^ spikes. In these experiments, the voltage-gated Na^+^ channel blocker QX-314 (2 mM) was included in the recording pipette to avoid the possibility of generating action potentials, as described (Nunez et al., 2012; Polsky et al., 2009).

In all experiments, basal PSP were recorded for 10-15 min before any drug application. BK blocker paxilline (1 µM) and NMDAR antagonist AP5 (100 µM) were applied in the bath. The parameters studied were PSP peak amplitude, area, and rise and decay times (Figures 5B and 5D).

##### Spike-timing-dependent plasticity

Induction of spike-timing-dependent plasticity (STDP) long-term potentiation (t-LTP) was achieved by pairing pre- and postsynaptic action potentials 10 ms away. Presynaptic action potentials were evoked by electrical stimulation of basal afferent inputs as described above and are recorded as PSP responses (Figure 6A). Postsynaptic action potentials were elicited by a brief current injection through the recording pipette (5 ms, 200-400 pA) as aforementioned. Basal PSP were recorded for 10-15 min before the pre-post associations were induced (30, 50, and 90 pairings were studied). After that, PSP were recorded for another 50-60 min. Amplitude of PSP 5 min before (-5 to 0 min interval) and 50 min after (45 to 50 min interval) the t-LTP induction protocol were measured to compare the extent of the PSP potentiation. For a pharmacological characterization of the t-LTP, BK blocker paxilline (500 nM) was applied intracellularly (through the recording pipette) (Figure 6B, right panel), whereas the NMDAR antagonist AP5 (100 µM) was applied in the bath (Figure Supp. 2B).

In some experiments, presynaptic stimulation of apical afferent inputs was paired with evoked postsynaptic action potentials to compare basal vs. apical t-LTP in BC-L5PN (Figure 6D). In these experiments, presynaptic electrical stimulation was performed in the limit between layers 1 and 2 (Figure 6C).

##### BK current recordings

BK currents from BC-L5PN were recorded as the paxilline-sensitive currents using the voltage-clamp mode in the whole-cell configuration of the patch-clamp technique in normal ACSF (2 MgSO_4_ and 2 CaCl_2_) supplemented with TTX (1 µM). Patch pipettes were filled with a modified recording solution (composition in mM: 123 KMeSO_3_, 9 NaCl, 9 HEPES, 0.9 EGTA, 14 Tris-phosphocreatine, 2 ATP-Mg, 2 ATP-Na, and 0.3 GTP-Tris; pH 7.3), as previously described (Whitt et al., 2018). I-V relationships were elicited from a holding potential of -70 mV, stepping from -110 to 90 mV for 150 ms in 20 mV increments (Figure Supp. 3). BK currents were isolated by current subtraction after bath application of the BK blocker paxilline (1 µM) and normalized to BC-L5PN capacitance (Whitt et al., 2018).

#### HEK293T cell line culture and transient ion channel transfection

Human embryonic kidney (HEK293T; ATCC #CRL-3216) cells were grown on 12-mm poly-lysine treated glass coverslips in Dulbecco’s Modified Eagle Medium supplemented with 10% fetal bovine serum and 1% penicillin-streptomycin. Cells were maintained in an incubator at 37°C in a 5% carbon dioxide atmosphere. Transient transfection of different combinations of cDNA plasmids encoding for BK alpha subunit (pBNJ-hsloTAG) and GluN1 (pEYFP-NR1a), GluN2A (pEGFP-NR2A), and GluN2B (pEGFP-NR2B) NMDAR subunits was performed using jetPRIME reagent (Polyplus Transfection). pBNJ-hsloTAG plasmid was generated in the lab (Giraldez et al., 2005) and pEYFP-NR1a (Addgene plasmid #17928), pEGFP-NR2A (Addgene plasmid #17924), and pEGFP-NR2B (Addgene plasmid #17925) plasmids were a gift from Stefano Vicini (Luo et al., 2002).

#### Electrophysiology in HEK293T cells

Poly-lysine treated glass coverslips containing the HEK293T cells expressing different combinations of NMDAR and BK channels were placed in a recording chamber mounted onto an inverted Nikon Eclipse Ti-U microscope (Nikon) equipped with 20x/40x objectives. Electrophysiological recordings were carried out 36–48 h post-transfection using the inside-out configuration of the patch-clamp technique (Figure 3C to 3H) (Hamill et al., 1981; Kshatri et al., 2018b). All experiments were performed at room temperature (22-24°C).

##### Inside-out BK current recordings in symmetrical K^+^ solutions

Patch pipettes were fabricated from GC150F-10 borosilicate glass capillaries as aforementioned and then fire-polished with a MF-200 microforge (World Precision Instruments) to obtain a tip resistance of 2–5 MΩ when filled with the recording solution (composition in mM: 80 KMeSO_3_, 60 N-methylglucamine-MeSO_3_, 20 HEPES, 2 KCl, and 2 CaCl_2_; pH 7.4). Recording solution was supplemented with NMDA (200 µM) and glycine (10 µM) to ensure NMDAR activation during the experiment, as previously described (Borschel et al., 2011; Xiong et al., 1998). Cells were superfused with a bath solution, containing (in mM): 80 KMeSO_3_, 60 N-methylglucamine-MeSO_3_, 20 HEPES, 2 KCl, and 1 HEDTA (pH 7.4). CaCl_2_ was added to obtain the desired free Ca^2+^ concentration. For Ca^2+^ solutions containing 100 μM Ca^2+^, no HEDTA chelator was added. The total amount of CaCl_2_ needed to obtain the desired Ca^2+^ concentration was calculated using the MaxChelator program (MAXC; (Bers et al., 2010)) and free Ca^2+^ was confirmed using a Ca^2+^-sensitive electrode (ThermoLab Systems). Stimulus generation and data acquisition were controlled and analyzed with the pClamp software package. The currents were amplified using an Axopatch-200B amplifier and digitized using a Digidata 1550A interface board (Molecular Devices). Subsequently, the data were acquired at 100 kHz and lowpass-filtered at 5 kHz. BK currents were elicited from a holding potential of -60 mV, stepping from -100 to 200 mV for 25 ms in 20 mV increments and then repolarizing to -80 mV for 100 ms to record the “tail currents”.

Conductance-voltage (G-V) curves were generated from tail current amplitudes normalized to the maximum amplitude obtained in 100 μM Ca^2+^. A Boltzmann equation was fitted to the data according to the following equation:

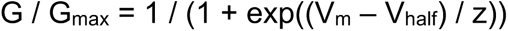

where V_half_ is the voltage of half-maximum activation, z is the slope of the curve, V_m_ is the test potential, and G_max_ is the maximal conductance (Figures 3C and 3D).

##### Inside-out BK current recordings in slices recording solutions

In some experiments (Figures 3F to 3H), inside-out recordings were performed using slightly modified brain slices solutions (see below). Briefly, the recording solution contained normal ACSF supplemented with NMDA (200 µM) and glycine (10 µM) to ensure NMDAR activation during the experiment. The bath solution contained (in mM): 135 KMeSO_4_, 10 KCl, 10 HEPES, 5 NaCl, 2.5 ATP-Mg, and 0.3 GTP-Na (pH 7.3), and was supplemented with EGTA (10 mM) to obtain a Ca^2+^-free solution. BK currents were elicited from a holding potential of -60 mV, stepping from -100 to 200 mV for 100 ms in 20 mV increments.

G-V curves were generated from I-V relationships, calculating the relative G for each test potential according to the following equation:

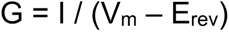

where I is the current amplitude at the end of the depolarizing pulse for each test potential (V_m_) and E_rev_ is the reversal potential for K^+^ with the solutions used. Individual G values were then normalized to maximal conductance (G_max_) and plotted as a function of V_m_ to obtain the G-V curves. A Boltzmann equation was fitted to the resultant values, as described before, and the V_half_ were obtained.

##### Proximity Ligation Assay

Proximity ligation assay (PLA) was performed using a commercially available kit (DuoLink; Sigma-Aldrich). HEK293T cells expressing different combinations of NMDAR and BK channels were fixed with 4% paraformaldehyde for 20 min, permeabilized, and then blocked for 1 h at 37°C to avoid non-specific binding of antibodies. BK channel was detected using a rabbit polyclonal anti-Maxi K^+^ channel alpha subunit primary antibody (1:200, #ab219072; Abcam). GluN1, GluN2A, and GluN2B subunits of NMDAR were detected using a goat polyclonal anti-NMDAR1 (1:200, #NB100-41105; Novus Biologicals) and mouse monoclonal anti-NMDAε1 (1:200, #sc-515148; Santa Cruz Biotechnology) and anti-NMDAε2 (1:200, #sc- 365597; Santa Cruz Biotechnology) primary antibodies, respectively. Secondary antibodies conjugated with oligonucleotides were supplied with the PLA DuoLink kit. Controls consisted of untransfected HEK293T cells or cells where only the BK alpha subunit or any of the NMDAR subunits alone were expressed (e.g., GluN1, as shown in Figure 3A). Image acquisition was performed using a Leica TS8 inverted confocal microscope (Leica Biosystems) and analysis and quantification were performed using Duolink Image Tool software (Sigma-Aldrich). Additional analysis and representation were performed using Fiji (Schindelin et al., 2012). A fluorescent dot corresponded with one protein-protein interaction, that is, where the proteins of interest are close enough (less than 40 nm) to be detected by the PLA technique (Alam, 2018). Results are expressed as the number of fluorescent dots (PLA signals) per cell area (in µm^2^). Four independent experiments were performed for each condition.

#### Drugs and Reagents

D-AP5 (D-2-amino-5-phosphonovalerate, AP5), glycine, ifenprodil (IFEN), N-methyl-D-aspartate (NMDA), paxilline (PAX), QX-314, and tetrodotoxin (TTX) were purchased from Tocris. EGTA and HEDTA were purchased from Sigma-Aldrich. BAPTA and ZnCl_2_ were purchased from Abcam and Merck, respectively. See Key Resources Table for complete references of all the drugs used in this study. Drugs were dissolved in the adequate solvent (DMSO or water) and prepared as 1000-10000x stock solutions of the desired final concentration. Subsequent dilution in ACSF (bath application) or pipette recording solution was performed to obtain the final concentration used in the experiments.

#### Statistical analysis

Data were analyzed with GraphPad Prism8 (GraphPad). Data are shown as mean ± SEM or as individual values (symbols) plus median and 25^th^-75^th^ percentile (boxes) values. A two-tailed t-test was used to analyze data from the same neuron measured twice (before/after) (e.g., pharmacological characterization in Figure 1E or t-LTP in Figures 6 and 7). A two-tailed unpaired t-test (Gaussian distribution) or a Mann-Whitney U test (non-Gaussian distribution) were used to analyze A-type vs. B-type BC-L5PN electrophysiological data. Kruskal-Wallis test was used to analyze PLA data. Statistical significance is stated as *p<0.05, **p<0.01, or ***p<0.001. Statistical details related to main and supplementary figures are specified in Table Supp. 1.

