## Supplementary text and Figures Gomez et al for "NMDA receptor–BK channel coupling regulates synaptic plasticity in the barrel cortex"

SUPPLEMENTAL INFORMATION

Supplementary Figures 1 to 5

Supplementary Table 1

### NMDA receptor–BK channel coupling regulates synaptic plasticity in the barrel cortex

**Ricardo Gómez, Laura E. Maglio, Alberto J. Gonzalez-Hernandez, Belinda Rivero-Perez[1], David Bartolomé-Martín, and Teresa Giraldez**

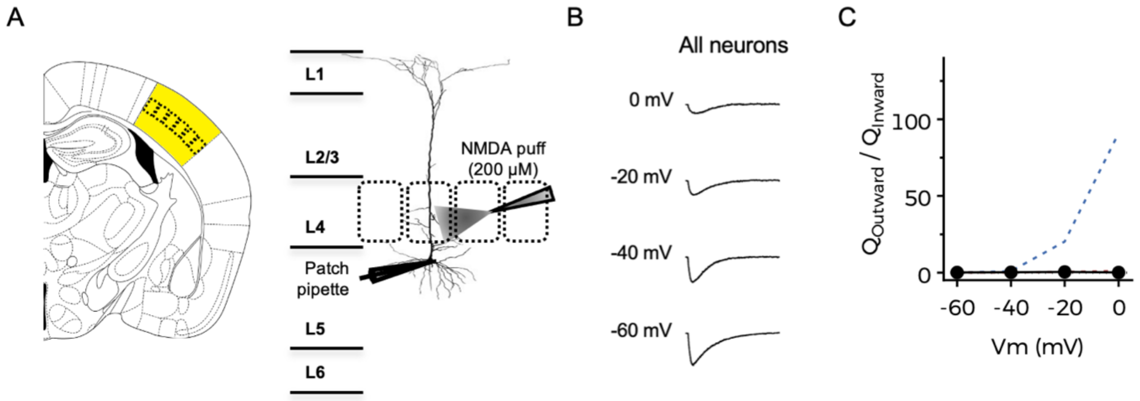


**Supplementary Figure 1 (related to Figure 1). NMDAR-dependent outward currents are absent in oblique dendrites and the initial segment of the apical dendrite**

**(A)** Left, general representation view of a mouse brain slice with the barrel cortex area highlighted in yellow. Right, schematic representation of the experimental design. **(B)** Representative current traces obtained at the indicated holding potentials after NMDA application. Scale bars represent 10 s and 200 pA. **(C)** Average Q-V relationships. Data points represent mean ± SEM; n=12. Dashed lines represent data from Figure 1C for a better comparison.

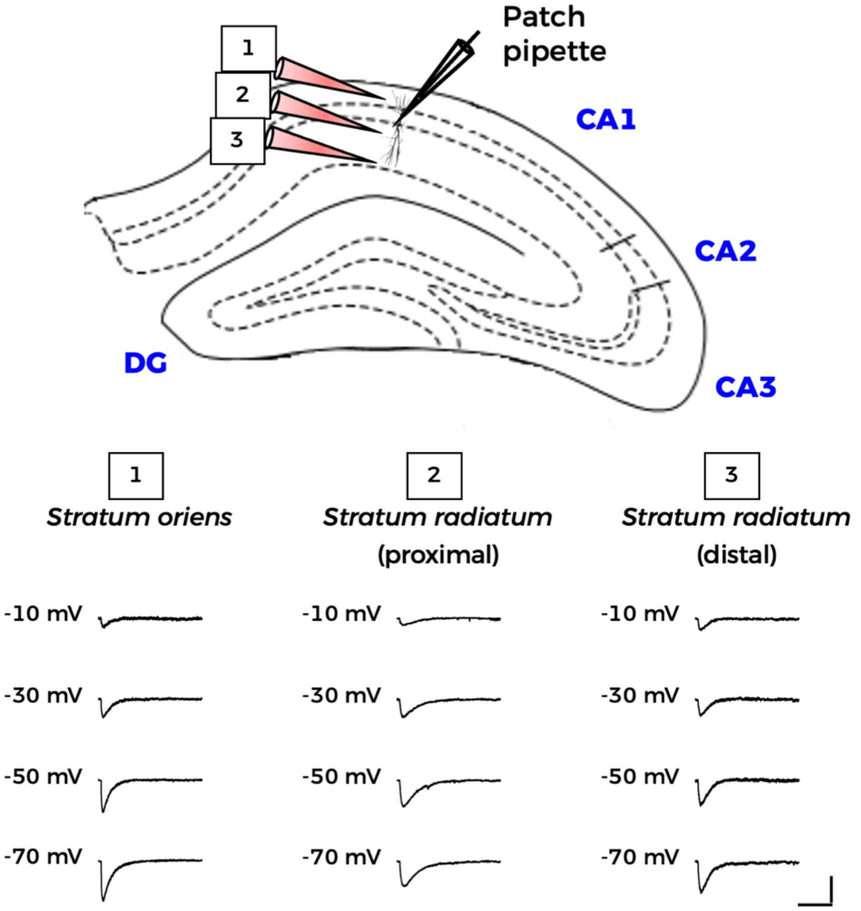


**Supplementary Figure 2. NMDA-evoked currents in dendrites from hippocampal CA1 pyramidal neurons**

Representative current traces obtained at the indicated holding potentials after NMDA application at different dendrite locations of hippocampal CA1 pyramidal neurons. No outward currents were observed in any case. Scale bars represent 10 s and 200 pA.

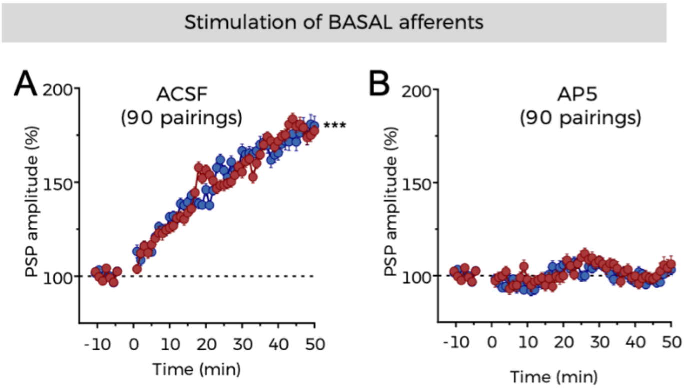


**Supplementary Figure 3 (related to Figure 6). NMDAR activation is mandatory for t-LTP induction**

**(A)** Time course of t-LTP development over time in A-type (red) and B-type neurons (blue) in control conditions (ACSF), following the experimental design depicted in Figure 6A. **(B)** Same experiments as in panel **A** were performed in the presence of 100 μM AP5. Data points represent mean ± SEM. A-type (ACSF): n=5; B-type (ACSF): n=6; A-type (AP5): n=5; B-type (AP5) n=4. Data in panel **A** are the same as Figure 7C (right panel) and are shown here for a better comparison. In **A**, ***p<0.001 (t-LTP vs. basal conditions). See also Table Supp. 1.

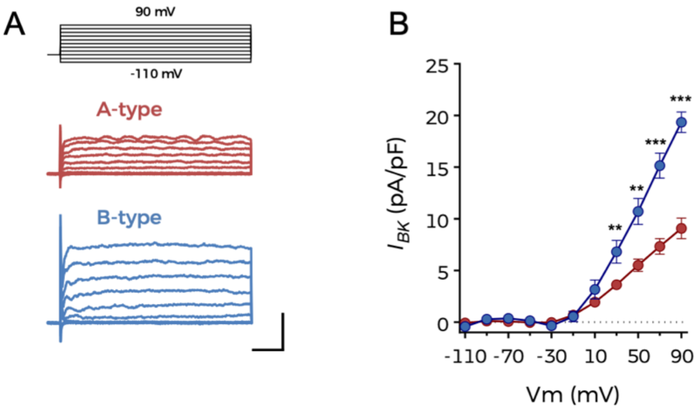


**Supplementary Figure 4 (related to Figure 6). BK channels are present in the plasma membrane of both types of BC-L5PN**

**(A)** Representative BK current traces obtained from an A-type (red) and a B-type neuron (blue) using the voltage protocol shown at the top and obtained as the paxilline-sensitive currents. **(B)** BK current density for A-type (red; n=8) and B-type (blue; n=4) neurons as a function of different membrane potentials. Data points represent mean ± SEM. In **B**, **p<0.01 and ***p<0.001 (B-type vs. A-type). See also Table Supp. 1.

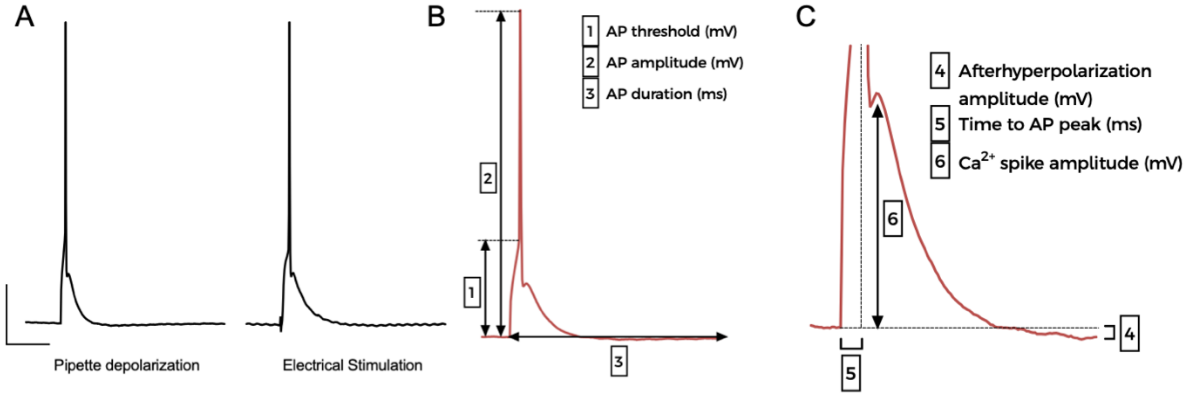


**Supplementary Figure 5. Single action potential characteristics and measurements**

**(A)** Representative single action potentials recorded from the same A-type BC-L5PN, evoked through pipette depolarization (left) or after electrical stimulation of basal afferent synaptic inputs (right). **(B)** Schematic description of the single action potential parameters determination summarized in Figure 4: threshold, amplitude, and total duration. **(C)** Determination of single action potential parameters summarized in Figure 4.

**Supplementary Table 1 (related to Statistical analysis). Summary of statistical significance and tests used in each figure** (comparisons reaching statistical significance are highlighted in light yellow)

| Figure 1. NMDAR activation opens BK channels in BC-L5PN basal dendrites | | | | |
|---|---|---|---|---|
| **PANEL** | **STATISTIC** | **CONDITION** | **P VALUE** | **OUTPUT** |
| Fig. 1C<br><br>Qoutward/ Qinward | Unpaired t-test (two-tailed) | -60 mV (B vs. A) | 0.1851 | Not significant |
| | | -40 mV (B vs. A) | 0.0990 | Not significant |
| | | -20 mV (B vs. A) | 0.0014 | ** |
| | | 0 mV (B vs. A) | 0.0041 | ** |
| Fig. 1D<br><br>NMDAR Current | Unpaired t-test (two-tailed) | -60 mV (B vs. A) | 0.9994 | Not significant |
| | | -40 mV (B vs. A) | 0.6021 | Not significant |
| | | -20 mV (B vs. A) | 0.9175 | Not significant |
| | | 0 mV (B vs. A) | 0.7962 | Not significant |
| Fig. 1D<br><br>NMDAR Charge | Unpaired t-test (two-tailed) | -60 mV (B vs. A) | 0.9997 | Not significant |
| | | -40 mV (B vs. A) | 0.7394 | Not significant |
| | | -20 mV (B vs. A) | 0.0449 | * |
| | | 0 mV (B vs. A) | 0.0435 | * |
| Fig. 1E (top)<br><br>Outward component | Paired t-test (two-tailed) | ACSF vs. AP5 | <0.0001 | *** |
| | | ACSF vs. $Zn^{2+}$ | 0.0002 | *** |
| | | ACSF vs. $Zn^{2+}$+AP5 | <0.0001 | *** |
| | | $Zn^{2+}$ vs. $Zn^{2+}$+AP5 | 0.0003 | ### |
| | | ACSF vs. IFEN | <0.0001 | *** |
| | | ACSF vs. IFEN+AP5 | <0.0001 | *** |
| | | IFEN vs. IFEN+AP5 | 0.0057 | ## |
| | | ACSF vs. PAX | <0.0001 | *** |
| | | ACSF vs. PAX+AP5 | <0.0001 | *** |
| | | PAX vs. PAX+AP5 | 0.5845 | Not significant |
| Fig. 1E (bottom)<br><br>Inward component | Paired t-test (two-tailed) | ACSF vs. AP5 | <0.0001 | *** |
| | | ACSF vs. $Zn^{2+}$ | 0.0781 | Not significant |
| | | ACSF vs. $Zn^{2+}$+AP5 | 0.0010 | *** |
| | | $Zn^{2+}$ vs. $Zn^{2+}$+AP5 | 0.0274 | # |
| | | ACSF vs. IFEN | 0.0785 | Not significant |
| | | ACSF vs. IFEN+AP5 | 0.0040 | ** |
| | | IFEN vs. IFEN+AP5 | 0.0413 | # |
| | | ACSF vs. PAX | 0.0381 | * |
| | | ACSF vs. PAX+AP5 | <0.0001 | *** |
| | | PAX vs. PAX+AP5 | 0.0283 | # |

| Figure 2. NMDARs and BK channels are within functional proximity in B-type BC-L5PNs | | | | |
|---|---|---|---|---|
| **PANEL** | **STATISTIC** | **CONDITION** | **P VALUE** | **OUTPUT** |
| Fig. 2E<br><br>EGTA<br>15 mM | Unpaired t-test<br>(two-tailed,<br>multiple t-test) | -60 mV (B vs. A) | 0.5963 | Not significant |
| | | -40 m V (B vs. A) | 0.0541 | Not significant |
| | | -20 m V (B vs. A) | 0.0040 | ** |
| | | 0 m V (B vs. A) | 0.0310 | * |
| Fig. 2F<br><br>BAPTA<br>1 mM | Unpaired t-test<br>(two-tailed,<br>multiple t-test) | -60 mV (B vs. A) | 0.6245 | Not significant |
| | | -40 m V (B vs. A) | 0.0020 | ** |
| | | -20 m V (B vs. A) | 0.0095 | ** |
| | | 0 m V (B vs. A) | 0.0058 | ** |

| Figure 3. Both GluN2A- and GluN2B-containing NMDARs can functionally couple to BK channels | | | | |
|---|---|---|---|---|
| **PANEL** | **STATISTIC** | **CONDITION** | **P VALUE** | **OUTPUT** |
| Fig. 3B<br><br>PLA | Kruskal-Wallis test | BK+GluN1/GluN2A<br>vs.<br>UNTRANSF, BK, and GluN1 | <0.0001 | *** |
| | | BK+GluN1/GluN2B<br>vs.<br>UNTRANSF, BK, and GluN1 | <0.0001 | *** |
| Fig. 3E<br><br>$V_{half}$<br><br>Symm. $K^+$ | Mann-Whitney U test<br>(two-tailed) | BK+2A vs. BK (0 $Ca^{2+}$) | <0.0001 | *** |
| | | BK+2A vs. BK (1 $Ca^{2+}$) | <0.0001 | *** |
| | | BK+2A vs. BK (10 $Ca^{2+}$) | 0.6165 | Not significant |
| | | BK+2A vs. BK (100 $Ca^{2+}$) | 0.0032 | ** |
| | | BK+2B vs. BK (0 $Ca^{2+}$) | 0.0003 | *** |
| | | BK+2B vs. BK (1 $Ca^{2+}$) | 0.0003 | *** |
| | | BK+2B vs. BK (10 $Ca^{2+}$) | 0.5743 | Not significant |
| | | BK+2B vs. BK (100 $Ca^{2+}$) | 0.0023 | ** |
| | | BK+2A vs. BK+2B | 0.5368 | Not significant |
| Fig. 3H<br><br>$V_{half}$<br><br>Slices sol. | Mann-Whitney U test<br>(two-tailed) | BK+2A vs. BK (0 $Ca^{2+}$) | 0.0008 | *** |
| | | BK+2B vs. BK (0 $Ca^{2+}$) | 0.0005 | *** |

| Figure 4. A subpopulation of regular-spiking BC-L5PNs exhibit NMDAR–BK functional coupling | | | | |
|---|---|---|---|---|
| **PANEL** | **STATISTIC** | **CONDITION** | **P VALUE** | **OUTPUT** |
| Fig. 4D | Mann-Whitney U test<br>(two-tailed)<br><br>B-type vs. A-type | Resting membrane potential | 0.4595 | Not significant |
| Fig. 4E | | Input resistance | 0.2386 | Not significant |
| Fig. 4F | | Capacitance | 0.8766 | Not significant |
| Fig. 4G | | Action potential freq. (200 pA) | 0.4088 | Not significant |
| | | Action potential freq. (230 pA) | 0.6359 | Not significant |
| | | Action potential freq. (260 pA) | 0.8089 | Not significant |
| | | Action potential freq. (290 pA) | 0.8481 | Not significant |
| | | Action potential freq. (320 pA) | 0.8815 | Not significant |
| Fig. 4H | | Action potential threshold | 0.2285 | Not significant |
| Fig. 4I | | Action potential amplitude | 0.6334 | Not significant |
| Fig. 4J | | Action potential duration | <0.0001 | *** |
| Fig. 4K | | $I_{AHP}$ amplitude | <0.0001 | *** |

| Figure 5. BK-dependent inhibition of NMDARs reduces postsynaptic response amplitude | | | | |
|---|---|---|---|---|
| **PANEL** | **STATISTIC** | **CONDITION** | **P VALUE** | **OUTPUT** |
| Fig. 5B<br><br>PSP Amplitude | Paired t-test (two-tailed) | A-type: ACSF vs. PAX | 0.0631 | Not significant |
| | | A-type: ACSF vs. PAX+AP5 | 0.0004 | *** |
| | | A-type: PAX vs. PAX+AP5 | 0.0003 | *** |
| | | B-type: ACSF vs. PAX | 0.0021 | ** |
| | | B-type: ACSF vs. PAX+AP5 | 0.0027 | ** |
| | | B-type: PAX vs. PAX+AP5 | 0.0004 | *** |
| Fig. 5B<br><br>PSP Area | Paired t-test (two-tailed) | A-type: ACSF vs. PAX | 0.8007 | Not significant |
| | | A-type: ACSF vs. PAX+AP5 | 0.0018 | ** |
| | | A-type: PAX vs. PAX+AP5 | 0.0179 | * |
| | | B-type: ACSF vs. PAX | 0.0029 | ** |
| | | B-type: ACSF vs. PAX+AP5 | 0.0010 | ** |
| | | B-type: PAX vs. PAX+AP5 | 0.0021 | ** |
| Fig. 5B<br><br>Rise Time | Paired t-test (two-tailed) | A-type: ACSF vs. PAX | 0.7043 | Not significant |
| | | A-type: ACSF vs. PAX+AP5 | 0.0785 | Not significant |
| | | A-type: PAX vs. PAX+AP5 | 0.3442 | Not significant |
| | | B-type: ACSF vs. PAX | 0.0008 | *** |
| | | B-type: ACSF vs. PAX+AP5 | 0.0014 | ** |
| | | B-type: PAX vs. PAX+AP5 | <0.0001 | *** |
| Fig. 5B<br><br>Decay Time | Paired t-test (two-tailed) | A-type: ACSF vs. PAX | 0.4604 | Not significant |
| | | A-type: ACSF vs. PAX+AP5 | 0.0028 | ** |
| | | A-type: PAX vs. PAX+AP5 | 0.0076 | ** |
| | | B-type: ACSF vs. PAX | 0.0004 | *** |
| | | B-type: ACSF vs. PAX+AP5 | 0.0030 | ** |
| | | B-type: PAX vs. PAX+AP5 | <0.0001 | *** |
| Fig. 5D<br><br>PSP Amplitude | Paired t-test (two-tailed) | A-type: ACSF vs. PAX | 0.3446 | Not significant |
| | | A-type: ACSF vs. PAX+AP5 | 0.0011 | ** |
| | | A-type: PAX vs. PAX+AP5 | 0.0102 | * |
| | | B-type: ACSF vs. PAX | 0.0015 | ** |
| | | B-type: ACSF vs. PAX+AP5 | 0.0011 | ** |
| | | B-type: PAX vs. PAX+AP5 | 0.0003 | *** |
| Fig. 5D<br><br>PSP Area | Paired t-test (two-tailed) | A-type: ACSF vs. PAX | 0.1448 | Not significant |
| | | A-type: ACSF vs. PAX+AP5 | 0.0013 | ** |
| | | A-type: PAX vs. PAX+AP5 | 0.0158 | * |
| | | B-type: ACSF vs. PAX | 0.0092 | ** |
| | | B-type: ACSF vs. PAX+AP5 | 0.0073 | ** |
| | | B-type: PAX vs. PAX+AP5 | 0.0040 | ** |
| Fig. 5D<br><br>Rise Time | Paired t-test (two-tailed) | A-type: ACSF vs. PAX | 0.0518 | Not significant |
| | | A-type: ACSF vs. PAX+AP5 | 0.4969 | Not significant |
| | | A-type: PAX vs. PAX+AP5 | 0.8170 | Not significant |
| | | B-type: ACSF vs. PAX | 0.0303 | * |
| | | B-type: ACSF vs. PAX+AP5 | 0.5845 | Not significant |
| | | B-type: PAX vs. PAX+AP5 | 0.0237 | * |
| Fig. 5D<br><br>Decay Time | Paired t-test (two-tailed) | A-type: ACSF vs. PAX | 0.4425 | Not significant |
| | | A-type: ACSF vs. PAX+AP5 | 0.0124 | * |
| | | A-type: PAX vs. PAX+AP5 | 0.0648 | Not significant |
| | | B-type: ACSF vs. PAX | 0.0394 | * |
| | | B-type: ACSF vs. PAX+AP5 | 0.0339 | * |
| | | B-type: PAX vs. PAX+AP5 | 0.0047 | ** |

| Figure 6. NMDAR–BK coupling increases the threshold for induction of synaptic plasticity | | | | |
|---|---|---|---|---|
| **PANEL** | **STATISTIC** | **CONDITION** | **P VALUE** | **OUTPUT** |
| Fig. 6B<br><br>Basal afferents (ACSF) | Paired t-test (two-tailed) | A-type (30 p)<br>Basal vs. t-LTP | <0.0001 | *** |
| | | B-type (30 p)<br>Basal vs. t-LTP | 0.4115 | Not significant |
| | Unpaired t-test (two-tailed) | B-type vs. A-type (30 p)<br>t-LTP | <0.0001 | ### |
| Fig. 6B<br><br>Basal afferents ($PAX_{int}$) | Paired t-test (two-tailed) | A-type (30 p)<br>Basal vs. t-LTP | <0.0001 | *** |
| | | B-type (30 p)<br>Basal vs. t-LTP | <0.0001 | *** |
| | Unpaired t-test (two-tailed) | B-type vs. A-type (30 p)<br>t-LTP | 0.9336 | Not significant |
| Fig. 6D<br><br>Apical afferents (ACSF) | Paired t-test (two-tailed) | A-type (30 p)<br>Basal vs. t-LTP | 0.3306 | Not significant |
| | | A-type (90 p)<br>Basal vs. t-LTP | <0.0001 | *** |
| | | B-type (90 p)<br>Basal vs. t-LTP | <0.0001 | *** |
| | Unpaired t-test (two-tailed) | B-type vs. A-type (90 p)<br>t-LTP | 0.7896 | Not significant |

| Figure 7. A high number and frequency of pre-post pairings relieves BK-dependent NMDAR inhibition | | | | |
|---|---|---|---|---|
| **PANEL** | **STATISTIC** | **CONDITION** | **P VALUE** | **OUTPUT** |
| Fig. 7A<br><br>t-LTP 0.20 Hz | Paired t-test (two-tailed) | A-type (30 p)<br>Basal vs. t-LTP | <0.0001 | *** |
| | | B-type (30 p)<br>Basal vs. t-LTP | 0.4115 | Not significant |
| | | A-type (50 p)<br>Basal vs. t-LTP | <0.0001 | *** |
| | | B-type (50 p)<br>Basal vs. t-LTP | <0.0001 | *** |
| | | A-type (90 p)<br>Basal vs. t-LTP | <0.0001 | *** |
| | | B-type (90 p)<br>Basal vs. t-LTP | <0.0001 | *** |
| Fig. 7B<br><br>Summary 0.20 Hz | Unpaired t-test (two-tailed) | B-type vs. A-type (30 p)<br>t-LTP | <0.0001 | ### |
| | | B-type vs. A-type (50 p)<br>t-LTP | <0.0001 | ### |
| | | B-type vs. A-type (50 p)<br>t-LTP | <0.0001 | ### |
| Fig. 7C<br><br>t-LTP 0.33 Hz | Paired t-test (two-tailed) | A-type (30 p)<br>Basal vs. t-LTP | <0.0001 | *** |
| | | B-type (30 p)<br>Basal vs. t-LTP | <0.0001 | *** |
| | | A-type (50 p)<br>Basal vs. t-LTP | <0.0001 | *** |
| | | B-type (50 p)<br>Basal vs. t-LTP | <0.0001 | *** |
| | | A-type (90 p)<br>Basal vs. t-LTP | <0.0001 | *** |
| | | B-type (90 p)<br>Basal vs. t-LTP | <0.0001 | *** |

| Figure 7 (continuation). A high number and frequency of pre-post pairings relieves BK-dependent NMDAR inhibition | | | | |
|---|---|---|---|---|
| **PANEL** | **STATISTIC** | **CONDITION** | **P VALUE** | **OUTPUT** |
| Fig. 7D<br><br>Summary 0.33 Hz | Unpaired t-test (two-tailed) | B-type vs. A-type (30 p) t-LTP | <0.0001 | ### |
| | | B-type vs. A-type (50 p) t-LTP | <0.0001 | ### |
| | | B-type vs. A-type (90 p) t-LTP | 0.4680 | Not significant |
| **Supplementary Figure 3 (related to Figure 6). NMDAR activation is mandatory for t-LTP induction** | | | | |
| Fig. S2A<br><br>t-LTP 0.33 Hz (ACSF) | Paired t-test (two-tailed) | A-type (90 p) Basal vs. t-LTP | <0.0001 | *** |
| | | B-type (90 p) Basal vs. t-LTP | <0.0001 | *** |
| | Unpaired t-test (two-tailed) | B-type vs. A-type (90 p) t-LTP | 0.4680 | Not significant |
| Fig. S2B<br><br>t-LTP 0.33 Hz (AP5) | Paired t-test (two-tailed) | A-type (90 p) Basal vs. t-LTP | 0.1026 | Not significant |
| | | B-type (90 p) Basal vs. t-LTP | 0.0821 | Not significant |
| | Unpaired t-test (two-tailed) | B-type vs. A-type (90 p) t-LTP | 0.4210 | Not significant |
| **Supplementary Figure 4 (related to Figure 6). BK channels are present in the plasma membrane of both types of BC-L5PN** | | | | |
| Fig. S3B<br><br>BK current | Unpaired t-test (two-tailed) | -10 mV (B vs. A) | 0.8611 | Not significant |
| | | +10 mV (B vs. A) | 0.1025 | Not significant |
| | | +30 mV (B vs. A) | 0.0073 | ** |
| | | +50 mV (B vs. A) | 0.0012 | ** |
| | | +70 mV (B vs. A) | 0.0002 | *** |
| | | +90 mV (B vs. A) | <0.0001 | *** |
